# Target-agnostic discovery of Rett Syndrome therapeutics by coupling computational network analysis and CRISPR-enabled *in vivo* disease modeling

**DOI:** 10.1101/2022.03.20.485056

**Authors:** R. Novak, T. Lin, S. Kaushal, M. Sperry, F. Vigneault, E. Gardner, S. Loomba, K. Shcherbina, V. Keshari, A. Dinis, A. Vasan, V. Chandrasekhar, T. Takeda, J.R. Turner, M. Levin, D.E. Ingber

## Abstract

Many neurodevelopmental genetic disorders, such as Rett syndrome, are caused by a single gene mutation but trigger changes in expression and regulation of numerous other genes. This severely impair functions of multiple organs and organ systems beyond the central nervous system (CNS), adding to the challenge of developing broadly effective treatments based on a single drug target. This challenge is further complicated by the lack of sufficiently broad and biologically relevant drug screens, and the inherent complexity in identifying clinically relevant targets responsible for diverse phenotypes. Here, we combined human gene regulatory network-based computational drug prediction with *in vivo* screening in a population-level diversity, CRISPR-edited, *Xenopus laevis* tadpole model of Rett syndrome to carry out target-agnostic drug discovery, which rapidly led to the identification of the FDA-approved drug vorinostat as a potential repurposing candidate. Vorinostat broadly improved both CNS and non-CNS (e.g., gastrointestinal, respiratory, inflammatory) abnormalities in a pre-clinical mouse model of Rett syndrome. This is the first Rett syndrome treatment to demonstrate pre-clinical efficacy across multiple organ systems when dosed after the onset of symptoms, and network analysis revealed a putative therapeutic mechanism for its cross-organ normalizing effects based on its impact on acetylation metabolism and post-translational modifications of microtubules. Although traditionally considered an inhibitor of histone deacetylases (HDAC), vorinostat unexpectedly restored protein acetylation across both hypo- and hyperacetylated tissues, suggesting non-HDAC-mediated therapeutic mechanisms supported by proteomic analysis.

## INTRODUCTION

Rett Syndrome (Rett) is a neurodevelopmental disorder that is also known to clinically impact multiple organs and systems in the body. It is primarily caused by mutations of the gene encoding methyl-CpG binding protein (MeCP2), a ubiquitous nuclear protein that acts as a transcriptional repressor and activator affecting hundreds of genes across the genome (Chahrour and Zoghbi, 2007). Rett is characterized by a high degree of cellular heterogeneity, with a mixture of wild type and MeCP2 mutant cells within the tissues of the same individual (Renthal et al., 2018) that leads to subtle but widespread gene misregulation (Gabel et al., 2015). Clinical criteria and research on Rett have primarily focused on neuromotor, social, and cognitive impairments and other CNS symptoms (e.g., seizures, repetitive purposeless movements, regression, microcephaly, loss of speech, autistic features)(Colvin et al., 2003; Kerr et al., 2001). However, these patients also commonly exhibit multiple clinical symptoms outside of the CNS, including respiratory dysfunction, gastrointestinal issues, systemic inflammation, disrupted metabolism, circadian rhythm and sleep disturbances, abnormal bone density, kidney function, nociception derangement, which are a major source of morbidity that contribute to a shortened lifespan (Hagberg, 2002; Vashi and Justice, 2019). It is important to note that although Rett is considered a neurodevelopmental disorder and therapeutics development has focused on CNS-related endpoints, these system-wide symptoms are often the most disturbing to patients and families, and thus there is a need to ameliorate symptoms of this disease across the entire body.

Monogenic diseases are theoretically amenable to gene therapy treatment, and Rett has been shown to be reversible in a mouse model of X chromosome reactivation (Przanowski et al., 2018) and post-natal expression of MeCP2 (Giacometti et al., 2007; Guy et al., 2007). However, even if current challenges around delivery efficacy and safety could be overcome, Rett and other X-linked disorders require gene dosage compensation to avoid toxicity due to overexpression of the corrected gene since even a modest 20% deviation of MeCP2 expression levels can lead to Rett symptoms (Collins et al., 2004; Luikenhuis et al., 2004). Therefore, pharmaceutical treatments for Rett and other neurodevelopmental disorders are still needed. Trofinetide, a synthetic IGF-1 C-terminal tripeptide activating BDNF expression, has shown initial promise in phase II clinical trials and it progressed to phase III trials but demonstrated only moderate efficacy (Glaze et al., 2017). Sarizotan, a 5-HT1A and D2 receptor agonist with promising efficacy in mouse models of Rett (Abdala et al., 2014), was recently terminated after clinical trial results failed to demonstrate efficacy (NCT02790034). Esketamine, an N-methyl D-aspartate receptor (NMDAR) agonist, is also being evaluated in the clinic (NCT03633058) due to its reduction of neuroexcitatory glutamate. But these and other therapeutic paths for Rett pursued thus far target only a subset of patients’ CNS symptoms and have not shown any effects on extra-CNS morbidity.

Here we present an integrated approach combining a novel gene expression network computational model to predict existing drugs and screening in an in vivo phenotypically diverse model of Rett syndrome generated in *Xenopus* tadpoles to assess whole-body efficacy. Our results led to the identification of vorinostat, which showed significant efficacy in a mouse Rett syndrome model even when initiating treatment after the onset of symptoms, demonstrating the utility of the target-agnostic platform for predicting drugs to restore multi-organ system function in Rett syndrome and potentially other complex diseases.

## METHODS

### *X. laevis* care and rapid generation of Rett syndrome tadpole models

*Xenopus* embryos and tadpoles were housed at 18°C with a 12/12 h light/dark cycle in 0.1X Marc’s Modified Ringer’s (MMR) medium. Tadpoles were fed 3x/week with sera Micron Nature fry food. All animal experiments and procedures were reviewed and approved by the Harvard Medical School (HMS) Institutional Animal Care and Use Committee regulations. Xenbase (Nenni et al., 2019) (www.xenbase.org) was used to search for MeCP2 homolog in *Xenopus laevis* (*Xenopus laevis* J-strain 9.2 version). MeCP2.L (XB-GENE-17346751), MeCP2.S (XB-GENE-494744), found on Chromosome 8L and 8S, show 75% and 70% sequence identity with human MeCP2, respectively. An important biological distinction is that while human MeCP2 is found on the X chromosome, Chromosome 8 is not part of the sex determination alleles of homomorphic chromosome pairs in *X. laevis*. Cas9 target sites were selected using CHOPCHOP (Labun et al., 2016) on the *X. laevis* J-strain 9.2 version, which facilitated the selection of guide RNA with no predicted off-targets. We used the default settings to generate a list of target sequences. We selected 4 targets from this list for each of the L and S forms. This selection was based primarily on the ranking provided by CHOPCHOP and as a function of their location on MeCP2 to target exons coding for the methyl-CpG-binding domain (MBD, exons 2 and 3) and Transcriptional repression domain (TRD, exon 3). The 8 sgRNA sequences are presented in **Supplementary Table 9** and were synthesized as modified sgRNA by Synthego Inc.

The sgRNA were resuspended to 100 μM in 0.1X Tris EDTA (pH 8.0). Then, an equimolar sgRNA mix was made using 2 μl of each sgRNA, for a total of 16 ul. The Cas9 RNP complex was formed by mixing 75 pmol of the sgRNA mix with 75 pmol of Cas9 in annealing buffer (5 mM HEPES, 50 mM KCl, pH 7.5), for a total volume of 100 μL, then incubated at 37°C for 10 minutes. The RNP was kept frozen at -20°C until the day of the injection. To generate MeCP2 knockdown models of Rett syndrome, *Xenopus* embryos were fertilized and maintained at 14°C until the 4-cell stage. Each cell was injected with ∼2 nL RNP per injection resulting in a final amount of 1.5 fmol of RNP per injection per cell, and we did not see any adverse effect when controlling for Cas9 or sgRNA at that amount. Note that the 1.5 fmol of ribonucleoprotein (RNP) used per injection is two to five time less than what has been commonly used in previously published studies for *Xenopus* (Aslan et al., 2017) as we found that higher amounts of sgRNA had adverse effects (not shown). Following injections, embryos were kept in 0.1X MMR at 18°C as described above, for the 18 days duration of the experiment.

### PCR and fragment analysis to validate MeCP2 gene disruption

MeCP2 editing efficiency was measured using Indel Detection by Amplicon Analysis (Yang et al., 2015), which leverages PCR using a pair of primers and universal fluorescein labeled oligo (**Supplementary Table 10**).The primers were synthesized by IDT. The amplicons were purified (Zymo Research) and submitted for fragment analysis on an ABI 3730XL (performed by Genewiz) following the IDAA method described previously. ProfileIT IDAA analysis software (COBO Technologies) was used to analyze the fragments and visualize the distribution of indels and visualize out-of-frame indels (Lonowski et al., 2017). IDAA is a robust method with 1-bp resolution suited for the analysis of a tetraploid genome without the need for a deep sequencing approach.

We analyzed the indels of 3 target sites on MeCP2.S of 14 tadpoles presenting with seizures and 3 non-edited tadpoles. Using a window of +/- 100 bp from the target site, we observed varied editing efficiencies depending on the target site. The likely out-of-frame indels ratio averaged 47% for Xl-mecp2s-g01 on exon 3, with the lowest being 12% for a given tadpole and highest being 95%, 48% for Xl-mecp2s-g02 on exon 2 (min/max 20-84%), and ∼82% for Xl- mecp2s-g04 on exon 2 (min/max 7-91%). Overall, 98% of indels where between 1 to 30 bp, and 84% were deletions.

### Microarray analysis

Tadpoles were sacrificed at stage 50 of development and lysed using a QIAGEN TissueLyser II bead mill (30 Hz, 2 x 30 sec) in a 2 mL tube with a 2.4 mm steel beads (Omni) and 1 mL phosphate buffered saline buffer at 4°C. 250 mL of lysate was used for RNA extraction using a QIAGEN RNeasy mini kit. RNA samples were DNAse treated. *Xenopus laevis* Genome 2.0 Array (Affymetrix) microarrays were processed by Advanced Biomedical Laboratories (Cinnaminson, NJ). RNA samples were processed using a Nugen Ovation PICO WTA System V2 kit. The resulting cDNAs were purified using a Qiagen MinElute PCR Purification Kit following the modifications outlined in the Nugen protocol. The cDNAs were fragmented and labeled using a Nugen Encore Biotin Module. Hybridization solutions were prepared by combining the fragmented, biotin-labeled cDNAs with hybridization cocktail (Affymetrix Hybridization, Wash, and Stain Kit). The mixtures were incubated in a thermal cycler at 99 °C for 2 min followed by 45 °C for 5 min. then loaded on *Xenopus laevis* Genome 2.0 arrays and incubated for 16-20 hours at 45 °C and 60 rpm in an Affymetrix Hybridization Oven 645. Following hybridization, arrays were washed and stained on Affymetrix Fluidics Station 450s using the Affymetrix FS450_0001 protocol with the stains and buffers supplied in the Affymetrix Hybridization, Wash, and Stain Kit. The stained arrays were scanned at 532 nm using an Affymetrix GeneChip Scanner 3000.

### Transcriptomics Data Processing

Microarray datasets were RMA normalized. RNAseq gene counts were normalized using size factors similar to the normalization performed by the R package DESeq. Genes missing values were removed and duplicates were combined by taking the maximum across counts. Gene expression heatmaps and hierarchical clustering plots were generated for all groups using the R package limma (version 3.42.2) and Python package seaborn (version 0.9.0). Volcano plots were generated using the Python package bioinfokit (version 0.7).

### nemoCAD Drug Prediction Algorithm

In the present study, we developed the nemoCAD computational tool to predict drugs that would shift MeCP2-edited tadpoles to a control tadpole state based on transcriptome state for each condition. nemoCAD utilizes pre-computed interaction probabilities of drug-gene and gene-gene interactions and differential gene expression signatures of a disease state and appropriate control to identify compounds capable of changing a transcriptional signature indicative of one biological state to another state of interest (e.g., reverting a disease state to a healthy state).

The repurposing algorithm is first used to identify transcriptome-wide differential expression profiles between two biological states in the input transcriptomic dataset (experimental or published) and to define the target normalization signature, i.e., the subset of genes whose expression levels need to be reversed in order to revert one state to the other (e.g., normalize the diseased state). Pairwise analysis, that implicitly assumes genes are expressed independently of one another, is carried out based on the comparison between gene expression profiles of >19,800 compounds found in the LINCS database release v1 (Subramanian et al., 2017) and the target normalization signature. Multiple correlation statistics (e.g., Pearson correlation, cross-entropy) across all differentially expressed genes are calculated and a combination score (e.g., Pearson correlation divided by the cross-entropy) is computed.

Putative predictions that incorporate gene-gene dependencies are then generated separately based on Bayesian network analysis on a regulatory and drug-gene interaction network architecture defined using publicly available databases of gene-gene interactions based on single gene knockout datasets in human cells (KEGG, TRRUST) and reference transcriptional signatures of drugs (LINCS, CTD), as described below. By combining a directed unweighted network structure with interaction probabilities for the connecting network edges, the constructed network is a weighted directed graph consisting of all possible paths that connect at least 2 genes of interest from the relevant genes within the target transcriptomic normalization set and the drugs; it therefore encodes the entire region of influence of a given list of genes and the drugs that can reverse their gene expression profiles in the desired manner.

This network is then used as input for a message-passing algorithm (e.g. loopy belief propagation algorithm (Forbes, 2021; Pearl, 1982)) in which the marginal probability distributions of drugs being “on” given the expression state of every gene are computed using the joint probability distribution encoded in the . This network-based approach leads to a ranking of all the drugs based on the probability of them inducing the desired transcriptomic signature across all genes in the network, which is separate from the initial ranking based on correlation analysis described above. This method has the advantage of incorporating gene-gene dependencies, but its success relies on the quality of network structures curated in the source databases.

Ultimately, drugs are prioritized that have both high correlation with the desired transcriptional signature change, representing desired drug-gene interactions, and high probabilities of being able to reverse the diseased state predicted by the Bayesian network analysis, representing gene network-level drug effects that can counter network-level disease processes. nemoCAD also enables visualization of the architecture of the network and subnetworks, and thus provides insight into potential molecular targets that can be verified using additional experiments. nemoCAD also can be used to optimize treatment regiments by iteratively inputting animal omics data following drug treatment to further tune drug predictions and develop drug combinations that might synergize in unpredictable ways. The approach underlying nemoCAD avoids the needs for *a priori* target or pathway inputs and, in fact, enables their discovery.

### Transcriptomics Gene Network Construction

Gene regulatory networks were inferred from gene expression data using the Bioconductor package GEne Network Inference with Ensemble of trees (GENIE3) release 3.12 in RStudio (R version 3.6.2)(Huynh-Thu et al., 2010). GENIE3 infers a weighted, directed gene regulatory network from expression data by decomposing the prediction of a network between *k* genes into *k* different regression problems. For each individual regression analysis, the expression pattern of one of the genes (target gene) is predicted from the expression patterns of all the other genes (regulator genes) using ensembles of regression trees. GENIE3 produces an adjacency matrix representation of the network with *k* nodes in which each node represents a gene, and an edge directed from one gene *i* to another gene *j* indicates that gene *i* regulates the expression of gene *j*. Based on known mechanisms involved in Rett Syndrome (**Supplementary Table 1**), a subset of genes was selected as candidate regulators: BDNF, FCRL2, FMR1, MeCP2, NTRK2, PER1, PER2, and PUM1. These potential regulators were calculated over all available target genes in the transcriptomics datasets (10,934 genes) for MeCP2 knock down and cas9-injected control samples. Digraph objects were constructed from adjacency matrices and visualized in Matlab R2020a (Mathworks; Natick, MA) for each of the MeCP2 knock down and control groups. The strongest regulator gene-target gene relationships were highlighted by filtering each network at a consistent edge weight (e_w_) threshold (e_w_ = 0.3) prior to plotting.

### Cross-Species Transcriptomics Analysis

Gene networks involved in Rett Syndrome in human patients and MeCP2 knock down in laboratory animal models were compared using previously acquired transcriptomics datasets (**Supplementary Table 2**). Transcriptomics datasets were processed using the same methods described for *Xenopus* transcriptomics and normalized using a series of transformations. Since RNA-seq data typically have a wider dynamic range than microarray datasets, RNA-seq and microarray data was transformed into a common space (Sekhon et al., 2013). RNA-seq datasets were transformed using a hyperbolic sine function and the log_2_ transformation was applied to microarray data (Sekhon et al., 2013). Subsequently, a variance stabilizing transformation was applied to remove potential mean-variance relationships in each dataset (Russo et al., 2018). Common genes expressed and measured between datasets (926 genes) were considered for further analysis. Dimensional reduction was performed using Principle Component Analysis (PCA) and visualized using Uniform Manifold Approximation and Projection (UMAP) in Python with the help of sklearn and UMAP packages.

Commonalities in gene expression values across species were analyzed by plotting heatmaps of differential gene expression using the Python packages seaborn and matplotlib. All datasets were normalized as described above and log fold change for each gene was calculated by combining samples of a condition using geometric mean. Log fold changes for all species were combined and the genes (96 genes) with variance greater than the mean variance (in fold changes) were used to construct the heatmaps.

Transcriptomics gene networks were constructed using GENIE3 and the same methods previously described. To investigate potential changes across the entire gene network, regulator genes were *not* down selected. Adjacency matrices were visualized for each network and the distribution of edge weights in the network was assessed by histogram and boxplots in Matlab. Digraph objects were constructed from adjacency matrices and visualized in Matlab for the strongest connections in each network (e_w_ > 99.99% for each network). Resultant networks were combined across all datasets for each species and plotted with different colors designating each dataset. A cross-species network that includes the strongest network connections across all datasets was formed by combining each species-specific dataset and plotted with different colors designating each species. To identify the strongest regulator and target gene nodes in the resultant cross-species network, the in-degree and out-degree were calculated in Matlab.

The in-degree is defined as the number of edges with that node as the target and out-degree of a node is equal to the number of edges with that node as the source.

### Drug screening

Drugs for tadpole screening were purchased from Sigma-Aldrich (St. Louis, MO, USA) or MedChem Express (Monmouth Junction, NJ, USA) and dissolved in DMSO to a stock concentration of 100-1,000 μM, depending on the solubility. For screens, compounds were diluted to their final concentrations with a 0.1% DMSO concentration. Media with dosed drug were made fresh and exchanged every two days following feeding. Dosing began at stage 45 and lasted for 7 days, after which standard 0.1X MMR was used to evaluate drug washout effects. Free-swimming tadpoles were imaged in 60 mm dishes using a SONY Alpha a6100 camera with 16mm objective against an illuminated background. Tadpole behavior was scored manually by noting the most severe seizure-related phenotype described by Hewapathirane et al. (Hewapathirane et al., 2008) within a 20 min period.

Vorinostat and trofinetide for mouse studies were purchased from MedChem Express (Monmouth Junction, NJ, USA). For intraperitoneal injection, trofinetide was solubilized in DMSO, and vorinostat was solubilized in a method previously published(Basu et al., 2019; Hockly et al., 2003). 2-hydroxypropyl-β-cyclodextrin powder (HPβCD) was purchased from Acros Organics (ThermoFisher, Waltham, MA, USA). Vorinostat was dissolved in 100 mM HPβCD by boiling for 5 minutes. Mice were dosed intraperitoneally at 100mg/kg with Trofinetide and 50 mg/kg with vorinostat. For oral dosing, commercial instructions were followed to add solubilized vorinostat (100 mg/kg) to Medigel (purchased from ClearH20, Westbrook, ME, USA).

### *X. laevis* Tissue Processing

After stage 46 tadpoles were euthanized with 20x Tricaine, tadpoles were washed three times with PBS (-/-) and fixed in 4% Paraformaldehyde in PBS + Mg + EGTA (MEMFA) (Alfa Aesar, Tewksbury, MA, USA) for 2 hours at RT. For sections, *Xenopus* tadpoles were cryoprotected by sequential incubation of 10%, 20% and 30% sucrose in 1x PBS(-/-), only transferring to the subsequent concentration when the samples sunk to the bottom of the scintillation vial. Cryoprotected tadpoles were embedded in 5% agarose and were stored in - 80°C. For GI staining, the gastrointestinal tract was microdissected with a stereoscope before embedding. Embedding tissue were sectioned coronally at 20 μm thickness on a Cryostat (Leica, CM3050 S, Wetzlar, Germany) and mounted on superfrost plus slides (Thermo Fisher, Waltham, MA, USA). Slides were dehydrated for 30 minutes at 20°C, before storing at -80°C. For whole mount preparation, tadpoles were processed in a method previously published (Willsey, 2021).

### *In situ* hybridization

Following X. *laevis* tissue processing of cryosections, sample pretreatment, RNAscope Target Retrieval and the RNAscope Assay was conducted by the HMS Neurobiology Imaging Facility following manufacturer’s instructions. Xl.MeCP2 was custom designed by Advanced Cell Diagnostics. Rpl8 was used as a positive control (Xl-LOC108706872-C3, Cat No. 516501-C3, Advanced Cell Diagnostics, Hayward, CA) and DapB as a negative control (Cat No. 320758, Advanced Cell Diagnostics, Hayward, CA).

### Immunohistochemistry of *Xenopus laevis* sections and whole mounts

Following tissue processing, cryosections were blocked for 2 hours at room temperature and subsequently incubated with primary antibodies overnight at 4°C, including mouse anti-alpha acetylated tubulin (1:1000 dilution, Sigma Aldrich, Burlington, MA USA), mouse anti-beta-tubulin (1:1000 dilution, E7, Developmental Studies Hybridoma Bank, Iowa City, Iowa, USA), anti-MeCP2 (1:1000 dilution, Invitrogen, Waltham, MA USA), Hoechst (1:1000 dilution, ThermoFisher, Waltham, MA, USA), negative controls anti-mouse IgG (1:1000 dilution, Invitrogen, Waltham, MA, USA), and anti-rabbit (1:1000 dilution, Invitrogen, Waltham, MA USA), as well as FITC-conjugated isolectin B_4_ (1:1000 dilution, Thermo Fisher, Waltham, MA, USA).

Slides were washed twice with TRIS-buffered saline, with 1% Tween-20 (TBT) for 5 minutes each and incubated in the dark with their corresponding secondary antibody: Goat anti-rabbit Alexa 647 (1:1000 dilution, Abcam, Waltham, MA, USA) and Goat anti-mouse Alexa 594 (1:1000 dilution, Abcam, Waltham, MA, USA) for 1 hour at room temperature. Slides were washed three times with TBT for 5 minutes each on an orbital shaker and mounted in ProLong Gold Antifade Mountant (ThermoFisher, Waltham, MA, USA) with a #1 coverslip and sealed. Slides were stored in the dark at 4°C before image acquisition. Z-stacks were acquired at 1*μ*m to visualize brain and gastrointestinal sections.

*X. laevis* embryos were processed for whole mount immunofluorescence as described (Willsey, 2021). Fixed tadpoles were quenched for 1 hour in a lightbox in 5% formamide and 4% hydrogen peroxide in PBS, permeabilized for 1 hour in PBS with 0.1% Triton X-100 (PBT) and blocked for 1 hour in 10% CAS-Block in PBT (Life Technologies, Carlsbad, CA, USA). Primary antibodies were incubated overnight at 4°C, including mouse anti-alpha acetylated tubulin (1:700 dilution Sigma Aldrich, Burlington, MA USA), mouse anti-beta-tubulin (1:100 dilution, E7, Developmental Studies Hybridoma Bank, Iowa City, Iowa, USA)), Hoechst (1:100 dilution, ThermoFisher, Waltham, MA, USA) and anti-mouse IgG (1:100 dilution). Slides were washed in PBT, blocked in 10% CAS-Block for 30 minutes, and incubated with the corresponding secondary antibody for 2 hours at room temperature including Goat anti-mouse Alexa 546 (1:100 dilution, Thermo Fisher, Waltham, MA, USA) and Goat anti-rabbit 633 (1:100 dilution, ThermoFisher, Waltham, MA, USA). Tadpoles are washed with PBT for 1 hour and then washed in PBS for another hour before mounting in Vectashield (Vector Laboratories, Burlingame, CA, USA). Z-stacks were acquired at 1*μ*m for visualizing epidermal MCC’s and 3*μ*m for visualizing entire brain regions.

### Drug target discovery in *X. laevis* using thermal proteome profiling

#### Sample preparation

While we largely followed the protein integral solubility alteration (PISA) thermal proteome profiling protocol (Gaetani et al., 2019), we adapted the mammalian cell- and tissue-specific parameters to tadpoles by optimizing the method to tadpoles by reducing thermal treatment temperatures to account for the lower normal temperature range of *X. laevis*.

Unmodified *X. laevis* tadpoles (stage 47-50) were exposed to 25 μM vorinostat or 0.1 μM ivermectin for 2 h while freely swimming in 0.1x MMR medium. Following euthanasia using Tricaine, animals were washed with fresh MMR, cut into thirds (head, abdomen containing all visceral organs, and tail), and placed into individual Eppendorf tubes covered with MMR containing the same drug and concentration and incubated for 3 min at one of 9 temperatures spanning 30-60°C. 30°C was selected as the control temperature, where no appreciable thermal denaturation should take place, based on the maximum tolerable temperature for *X. laevis* (Ruthsatz et al., 2018). Following aspiration of remaining liquid, samples were flash frozen in liquid nitrogen.

Samples were further processed and analyzed by Phenoswitch Bioscence, Inc. (Sherbrooke, Québec, Canada). Thermally-treated samples were lysed in 100 µL PBS with 0.4% NP-40 with 4 cycles of freeze/thaw. Insoluble material was cleared by centrifugation (10 min, 10,000 G, 4°C). 11µL of supernatant for each of the 9 temperatures were then pooled in one tube and samples were centrifuged again at 13,000 RPM for 75 minutes, at 4°C to pellet precipitated proteins. 80 µL of the supernatant were reduced with 10 mM DTT for 15 min at 65°C and alkylated with 15 mM IAA and 30 min at room temperature in the dark. Proteins were precipitated with 8 volumes of ice-cold acetone and 1 volume of ice cold methanol overnight. Protein pellets were washed 3 times with 250 µl of ice-cold methanol and resuspended in 100 µL digestion buffer. Digestion was carried for 4 hours in 50 mM Tris ph 8 + 0.75 mM Urea + 1µg trypsin/LysC at 37°C with agitation. Another 1 µg of trypsin/LysC was added, and digestion was continued overnight. Peptides were purified by reversed phase SPE and analyzed by LC-MS/MS.

#### Mass spectrometry

Acquisition was performed with a TripleTOF 6600 (Sciex, Foster City, CA, USA) equipped with an electrospray interface with a 25 μm iD capillary and coupled to a Micro LC200 (Eksigent, Redwood City, CA, USA). Analyst TF 1.8 software was used to control the instrument. Acquisition was performed in data independent acquisition (DIA or SWATH) using gas phase fractionation (GPF1 from 350 m/z to 800 m/z and GPF2 from 800 m/z to 1250 m/z). The source voltage was set to 5.5 kV and maintained at 325oC, curtain gas was set at 45 psi, gas one at 25 psi and gas two at 25 psi. Separation was performed on a reversed phase Kinetex XB column 0.3 μm i.d., 2.6 μm particles, 150mm long (Phenomenex) which was maintained at 60°C. Samples were injected by loop overfilling into a 5 μL loop. For the 60 minutes LC gradient, the mobile phase consisted of the following solvent A (0.2% v/v formic acid and 3% DMSO v/v in water) and solvent B (0.2% v/v formic acid and 3% DMSO in EtOH) at a flow rate of 3 μL/min. Both GPF files were analyzed on a previously-generated 3D ion library using the SWATH 2.0 microapp from Peakview (Sciex, Foster City, CA, USA). Each GPF file was analyzed using 10 peptides per protein, 4 MS/MS transition per peptide, 12.5 min RT window and 25 ppm XIC width. The reported quantification for a protein is the sum of all the correctly integrated peptides (FDR<0.05) in both GPF files.

### Mouse care

Experiments were performed with MeCP2-null (*MeCP2^-/Y^*) (strain 003890) and age-matched WT male littermates (*MeCP2^+/Y^*), purchased from Jackson Laboratories (Bar Harbor, ME). All mice were housed in ventilated racks under specific-pathogen-free conditions at a room temperature in a 12/12 h light/dark cycle with food and water ad libitum. All animal experiments and procedures were reviewed and approved by the Harvard Medical School (HMS) Institutional Animal Care and Use Committee regulations. Every effort was made to minimize their suffering. Mice were assessed for disease severity and endpoint criteria on a daily basis by weighing and using the phenotypic scoring method developed by the Bird laboratory (Guy et al., 2007; Szczesna et al., 2014), which includes assessing mobility, gait, breathing, tremor, and general condition. Endpoint was determined by 20% body mass loss or a score of 2 in criteria D, E, or F. Dosing and Behavioral procedures began when mice reached 4 weeks of age.

### Behavioral Testing

#### Elevated Plus Maze (EPM)

The elevated plus maze (EPM) consisted of two open and two closed (34 cm long and 5 cm wide) arms extended out from a central platform 50 cm above the floor. The test was carried out in dim ambient lighting. The room of the experiment and its spatial cues were kept consistent and minimal. Mice were habituated in the test room for 30 minutes before the start of the test. Mice were placed near the center compartment of the maze, facing an open arm, and allowed to explore the apparatus for 5 minutes. After each test, the maze was cleaned thoroughly with 70% ethanol and left to dry before the start of the next test. A computer-assisted video-tracking system (EthoVision XT 14, Noldus, Leesburg, VA) was used to record the number of open and closed arm entries as well as the total time spent in open, closed, and center compartments. An increase in the percent time spent or entries into the open arms was used as a surrogate measure of anxiolytic-like behavior (Carobrez and Bertoglio, 2005).

#### Spatial Novelty Y-Maze (Y-maze)

The spatial novelty y-maze consisted of a Y-shaped maze with three arms at a 120° from each other as well as a removable blockade used to prevent access to one of the arms during the Habitutation phase. The test was carried out in dim ambient lighting. The room of the experiment and its spatial cues were kept consistent. One of the three arms was defined as the start arm, where the mice would be placed at the start of the experiment and was kept consistent throughout the entire study. The test consisted of a three-minute Habituation phase where one of the two non-start arms was blocked off, a two-minute intertrial interval (ITI) outside of the maze and a three-minute Test phase, where the previously blocked arm was exposed to the mouse. The arm that was blocked off during the Habituation phase was randomized and noted for each mouse.

Mice were kept in an adjacent room to not disturb the test. The test mouse was placed in a holding cage and brought to the test room with the maze and immediately placed into the Y- maze at the designated start arm and allowed to explore the start and familiar arm trial for three minutes. At the end of the habituation phase, the mouse was placed into the holding cage for 2 minutes ITI. During this time, the blockade was removed, and the maze was cleaned with an ammonia-based cleaner with a paper towel and left to dry to remove odor cues. The mice were then placed back in the start arm and left to explore the entire maze for three minutes. After the test trial, the mouse was transported back to its home cages and the maze was again wiped dry with an ammonia based cleaner. A computer-assisted video tracking system (EthoVision XT 14, Noldus, Leesburg, VA) was used to record the number of entries, time spent in each arm and as distance traveled in each arm during the Habituation and Test Phase. The percentage of time and distance spent in the novel, or previously blocked arm were used to assess spatial novelty seeking behavior (Dellu et al., 2000).

### Mouse Histology and Immunostaining

Mice were euthanized with CO_2_ inhalation, transcardially perfused with PBS (-/-) and 4% PFA, and tissues of interest were removed. Regions of the gastrointestinal tract and lung processed as was described previously(Morton and Snider, 2017; Nalle et al., 2019; Tsai et al., 2017). Paraffin blocks were sectioned into 15 *μ*m thickness. Mouse lung tissues were sectioned coronally and GI tissues were stained with hematoxylin and eosin (H&E) by the Beth Israel Deaconess Medical Center’s IHC Core Facility. The slides were also stained for rabbit anti- acetylated *α*-tubulin (ABclonal, ab179484, Woburn, MA, USA) mouse anti-CD-64 (R&D systems, AF3628, Minneapolis, MN, USA), chicken anti-*β*III-tubulin (Biolegend, 801202, San Diego, CA, USA) immunofluorescence by Beth Israel Deaconess Medical Center’s IHC Core Facility.

Mouse brains were fixed with 4% paraformaldehyde at 4°C overnight, cryoprotected in a sucrose gradient (10%, 20% and 30%), and subsequently embedded in OCT Tissue Tek. The embedded tissues were subsequently sectioned coronally at 30*μ*m and the free-floating sections were stained as previously described(Potts et al., 2020). The sections were blocked with 10% CAS Block in PBT for 1hr, stained with primary antibody (rabbit anti-iba-1, 1:1000, Wako, #019- 19741; mouse anti-NeuN, 1:1000, Abcam, #ab104224; Hoechst, 1:1000, Thermo Fisher, H1399) overnight at 4°C. After three 10-minute washes in PBS (-/-) on an orbital shaker, the sections were stained in secondary antibody (anti-rabbit Alexa 647 for iba1, anti-mouse Alexa 594 for NeuN) for 1 hour at room temperature. After three 10-minute washes on an orbital shaker, the sections were mounted on superfrost plus slides (ThermoFisher, Waltham, MA, USA) in ProLong Gold Antifade Mountant (ThermoFisher, Waltham, MA, USA) with a #1 coverslip and sealed. Z-stack images were acquired at 1*μ*m intervals using the 63x/ glycerol.

### Multiplex Chemokine Assay

After mouse euthanization, blood was collected by cardiac puncture using at 23G needle in ethylendiamine tetra-acetic acid (EDTA) treated tubes (Microvette 200 K3EDTA, CAT# 20.1288.100, Sarstedt, Numbrecht, Germany). To avoid hemolysis, the tubes were kept at room temperature and quickly processed for plasma. Tubes were centrifuged at 2000 x g for 10 minutes and the plasma was separated and stored in -80C until assayed. Chemokine levels were measured in mouse plasma using the Bio-Plex Pro Mouse Chemokine Panel 31-Plex (Cat# 12009159, Bio Rad, Hercules, CA, USA). Measurements were performed using the Bio-Plex 3D Suspension Array System (BioRad, Hercules, CA, USA), following the manufacturer’s instructions.

### Microscopy and Image Processing

H&E slides were imaged with 4x and 20x objectives with the Biotek Cytation 5. Immunofluorescent *Xenopus* whole mounts and *Xenopus* and mouse stained sections were imaged with the Leica SP5 X MP Inverted Confocal Microscope using 25x (NA 0.95) water objective and 63x (NA 1.3) Glycerol objectives. Sections were imaged in white light and diode laser scanning mode and whole mounts were imaged using multiphoton pulsed IR laser scanning. Serial scanning was used during acquisition to avoid bleed-through. Image settings were kept constant between groups, in terms of laser power, bit-depth, z-stack interval, resolution (pixels), aperture settings, and gain/offset. Images were post-processed and analyzed using ImageJ (Fiji, NIH).

#### MCC morphological analysis

2-photon images of X. laevis 25x and 63x MCC’s were maximum projected, background subtracted, and subsequently assessed for cilia orientation and cilia length, as previously described (Chien et al., 2018).

#### MCC Functional Assay

We visualized the cilia-driven flow of fluorescent microspheres over the surface of the xenopus tadpole by taking times series with the Leica SP5 x MP inverted confocal. Our methods were based on previous reports (Kulkarni et al., 2018) with some modifications. Stage 45 tadpoles were placed in a 24 well *μ*-Plate 14mm (ibidi, Gräfelfing, Germany), and underwent anesthesia by incubation with 1x tricaine before proceeding to image. One tadpole was placed per well for individual tracking. Fluorescent microspheres beads (Thermo Fisher, Waltham, MA) were added at a final dilution of 1:100. We chose the bead size of 3*μ*m, because during preliminary testing and as is reflected in the data, WT *X. laevis* tadpoles consistently was able to clear beads that were 3*μ*m diameter at this concentration. The Leica SP5 x MP laser scanning mode was used to visualize the fluorescent microspheres beads at an excitation of 520nm and emission of 560nm with the following settings: 512x512 pixels, bidirectional scanning mode, and at a scan speed of 800-1600Hz, with the 25x (NA 0.95) water objective, and over the time course of 5 minutes every 5s. Time series were acquired of in the olfactory and lateral region of the epidermis.

#### Quantification of bead displacement

The time series stacks were maximum intensity projected to visualize the overall bead flow. Intensity was measured as a function from the distance from the surface of the epidermis.

#### Mean Intensity Quantitation

To generate a ROI mask, channels acquired in one z-stack were split and the maximum intensity projection of a ‘control’ channel was taken. In the case for assessing *α*aT expression mouse GI tract and bronchiole, the *β*III-tubulin channel was used, and for assessing MeCP2 expression, the *α* -acetylated tubulin channel was used in the image stack. Subsequently, background noise was subtracted, and auto-thresholded, whose settings were held constant between groups. The ROIs generated from the control channel were used to quantify the maximum projected channel of interest, to obtain the mean intensity (integrated density per area). A minimum of 3 serial sections were analyzed per animal.

#### Counting Ib4+ cells

Confocal Z-stacks of *X. laevis* GI tract images were quantified for IB4+ cells, by taking the maximum intensity projection of the ib4+ channel. Following noise reduction, and auto-threshold, ib4+ cells were segmented and counted using the particle analysis function in the ImageJ software (NIH, Bethesda, MD).

#### Sholl Analysis

Sholl analysis was performed as previously published^47^. Confocal (at 1um intervals) Z-stacks were maximum intensity projected in the iba-1 channel, noise de-speckled, subsequently segmented using the auto-threshold function, and microglia were duplicated into 8-bit TIF files, in the field of view, ensuring that for each mouse at least 10 microglia were selected in the ipsilateral and contralateral region of the olfactory bulb. Using the Sholl analysis plugin in ImageJ (NIH, Bethesda, MD), traces and intersections were generated.

### Statistical analysis

Statistical analysis of biological results was performed using Graphpad Prism 9.3. For comparison of two conditions, Student’s t-test was used unless otherwise indicated. ANOVA with pairwise t-tests and Bonferroni correction or the Holm-Šídák tests was used to evaluate significance of multiple variables. P values <0.05 were considered significant (*, p < 0.05; ** p < 0.01; *** p < 0.001; **** p < 0.0001). Tadpole screens and mouse studies were conducted with 5-8 animals each and repeated, except for the mouse oral dosing study.

## RESULTS

### Modeling Rett syndrome genetic and phenotypic heterogeneity in *Xenopus laevis*

A major challenge to Rett therapeutics discovery lies in the inherent heterogeneity observed in patients stemming from gene expression variability due to the gene regulatory impact of distinct MeCP2 mutations, combined with tissue heterogeneity caused by mosaicism resulting from X chromosome inactivation. As a result, Rett patients exhibit a spectrum of disabilities. To establish an animal model of Rett that would incorporate the heterogeneity observed in patients, we used CRISPR to generate a mosaic knockdown of MeCP2 protein in *Xenopus laevis* tadpoles, which have been previously used to model neurodevelopmental diseases due to their evolutionary proximity to mammals and well-characterized neural development (Exner and Willsey, 2021; Pratt and Khakhalin, 2013).

Injection of *X. laevis* embryos at 4- or 8-cell stages using cas9 pre-complexed with sgRNA targeting 6 sites of the MeCP2 gene, including the regions corresponding to the DNA-binding domain, resulted in a tunable model of MeCP2 knockdown with a heterogeneous phenotype when assessed at the population level. Based on our target selection and injection strategy at the 4-cell stage, each individual animal was edited as a mosaic and presented varying degree of severity, without apparent toxicity from Cas9 or sgRNA. Indeed, all embryos developed to the swimming tadpole stages (Nieuwkoop-Faber stages 45-50) with no apparent loss of viability or morphological defect in earlier stages.

Importantly, swimming tadpoles with MeCP2 knock down (“Rett” tadpoles) exhibited a broad range of abnormal behavior compared to wild type vehicle-injected control tadpoles (“Control”) including darting motions and rapid repetitive swimming in tight circles (**Fig. 1a,b and Supplementary Movie 1**) reminiscent of the repetitive motions observed in human Rett patients as well as C-shaped or straight rigor (**Supplementary Movie 1**) that has been shown to correspond to seizures in prior *Xenopus* studies (Hewapathirane et al., 2008). The cas9 protein and sgRNA doses were titered to achieve between 5 and 95% gene editing efficiency as determined by qPCR without resulting in toxicity (**Supplementary Fig. 1a**). All tadpoles exhibited significant MeCP2 gene insertion/deletion polymorphisms (indels) at each of the mecp2 gene target sites (**Supplementary Fig. 1b**) and reduced MeCP2 RNA expression (**Fig. 1c**). MeCP2 protein levels were also heterogeneously reduced in the brain (**Fig. 1d,e**) and olfactory multiciliated cells (MCC) (**Fig. 1d,f**). This this editing approach resulted in an animal model of Rett with intra-subject and inter-subject diversity at molecular and phenotypic scales, which is similar to what is observed in humans.

**Figure 1:**
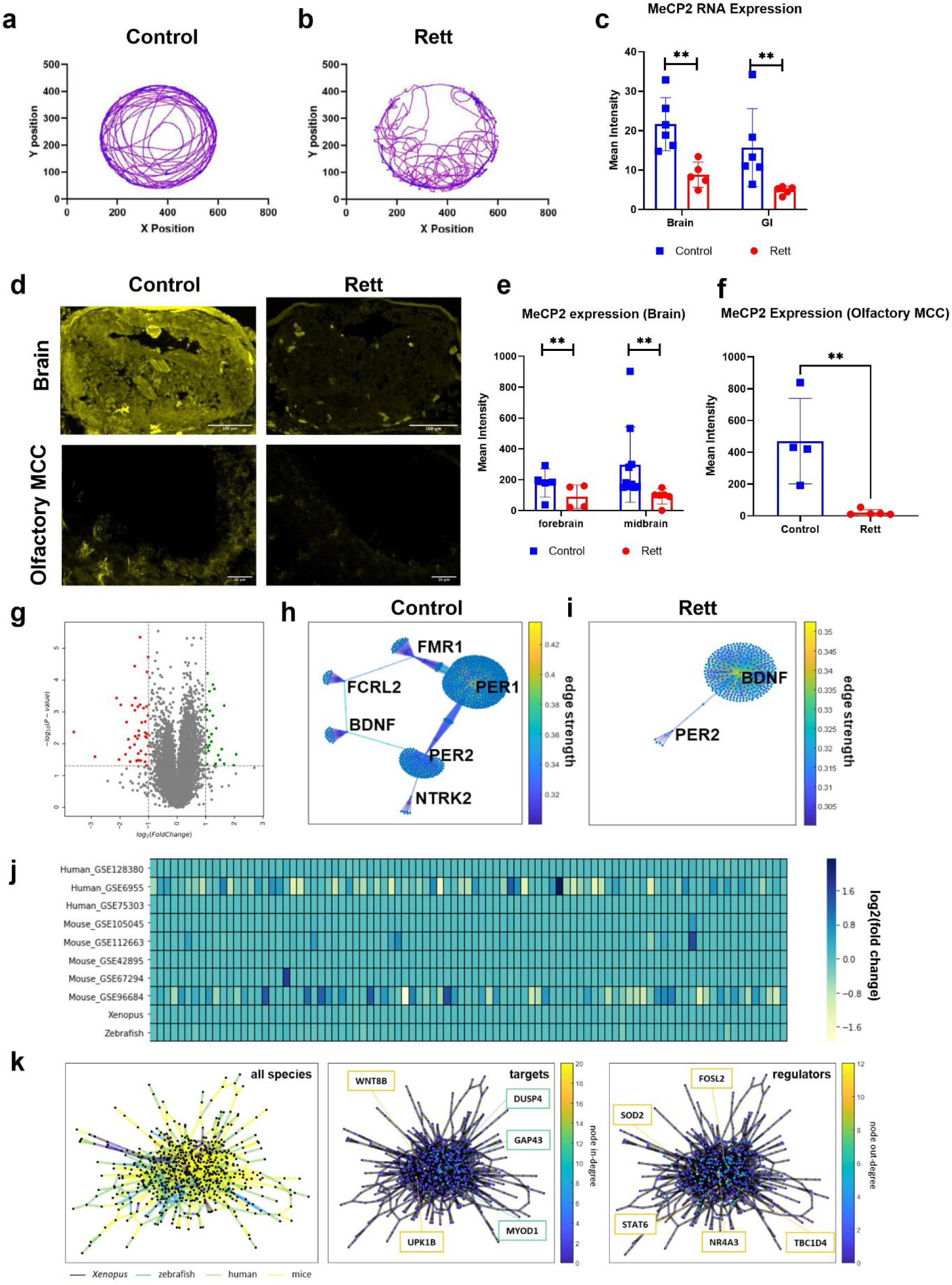
MeCP2 knockdown using CRISPR in *Xenopus laevis* tadpoles models Rett syndrome. a-b, Tadpole models of Rett syndrome exhibit distinct swimming behavior in 60 mm diameter dishes following MeCP2 knockdown (“Rett”) compared to mock-injected controls (“Control”). MeCP2 RNA expression in brain and gastrointestinal (GI) tract using RNAscope, **c**, and protein in brain and multi-ciliated olfactory cells using immunohistochemistry, **d-f**, are significantly reduced in Rett syndrome tadpole models while maintaining a large degree of heterogeneity spatially, **d**, and across tissues, **e-f**; scale bars, 100 μm in brain, 20 μm in olfactory multi-ciliated cells. **g**, Volcano plot of differentially expressed genes showing that only 70 genes are significantly up- or down-regulated following MeCP2 knockdown (p < 0.05, fold change >2, t-test with Bonferroni correction, N = 3). Gene regulatory networks for control, **h,** and Rett, **i,** tadpoles reveal large network rearrangements with an increase in BDNF centrality. **j**, Comparison against other Rett syndrome models and clinical samples indicates minimal differential gene expression across the 96-gene signature developed to classify Rett syndrome. **k**, Network-level comparison across species and tissues (left) and identified shared target (middle) and regulator (right) genes.

In line with prior studies in humans and other model organisms (Sanfeliu et al., 2019, p. 2), when we analyzed the transcriptome-wide effects of MeCP2 knockdown on the Rett phenotype in *Xenopus*, only subtle shifts in gene expression were detected with 70 out of 10,935 genes probed undergoing a significant change in gene expression (p_adjusted_<0.05) (**Fig. 1g, Supplementary Table 1**). Of the 70 differentially expressed genes, 37 are involved in metabolic processes, 9 operate within developmental processes, and 8 are part of signal transduction pathways (**Supplementary Fig. 2a**); this aligns with changes in genes that control metabolism or regulate neuronal processes (e.g., ion transport, nervous system development) in Rett patients (Renthal et al., 2018).

Interestingly, analysis of gene co-expression networks identified substantial reorganization from control animals (**Fig. 1h**) following MeCP2 knock down (**Fig. 1i**), characterized by a loss of strong connectivity among genes involved in the regulation of MeCP2 (FMR1 and TRKB receptor NTRK2 (Abuhatzira et al., 2007; Arsenault et al., 2021)), neuronal development (BDNF (Chang et al., 2006; Li and Pozzo-Miller, 2014)),, and circadian rhythm (PER1, PER2 (Martínez de Paz et al., 2015)) all of which are altered in Rett patients (Li and Pozzo-Miller, 2014; Miller et al., 2019). Notably, after MeCP2 knock down, BDNF strengthened as a network hub and exhibited greater co-expression connectivity amongst gene nodes (**Fig. 1i**).This was unexpected because although BDNF is involved in neuronal development, synaptic transmission, and plasticity, it declines with the onset of Rett-like neuropathological and behavioral phenotypes (Li and Pozzo-Miller, 2014). However, BDNF is a known target of repression by MeCP2 and our findings are consistent with prior work suggesting that downregulation of BDNF is a later and indirect outcome of MeCP2 deficiency (Chang et al., 2006; Li and Pozzo-Miller, 2014).

We then comprehensively compared the effects of MeCP2 knock down in *Xenopus* to changes observed in published transcriptomic data sets from Rett patients as well as MeCP2 knockout mouse and zebrafish models (**Supplementary Table 2**). After performing dimensionality reduction in this cross-species analysis, *K*-Means clustering applied to the principal components revealed 3 clusters, with mouse and human clustering together and zebrafish and *Xenopus* forming 2 independent clusters (**Supplementary Fig. 2b**). Importantly, diverse gene-expression abnormalities that occur across cell types in Rett syndrome could contribute to the differences observed across these datasets (Renthal et al., 2018). In this case, both the human and mouse transcriptomics were performed on brain tissues, whereas the zebrafish and *Xenopus* data are from whole organisms, which may account for the clustering patterns observed.

Despite evidence of species-specific clustering, a common gene signature comprised of 96 genes was identified across all species by searching for genes with a fold change variance greater than mean variance. However, the common genes within this transcriptomic signature are expressed at similar low fold change levels in 7 of the 9 datasets analyzed, with slightly greater variations in expression only observed in 1 out of 5 mouse datasets and 1 out of 3 human datasets (**Fig. 1j**). This finding provides additional evidence that mutation of MeCP2 produces widespread, yet subtle, perturbations in gene expression.

Using gene co-expression network analysis (Huynh-Thu et al., 2010), we also identified interconnected gene sub-networks that are conserved across species and tissues in the MeCP2 deficient condition (**Fig. 1k, Supplementary Tables 3 and 4**). Comparison across all species identified several shared major gene targets and regulators despite differences in species-specific network structure (**Supplementary Fig. 2c**). Interestingly, WNT8B, which is known to be modulated by the Rett-implicated gene Foxg1 (Aguiar et al., 2014), was found to be the most interconnected gene target within the MeCP2 cross-species network. The strongest gene regulators in this network, NR4A3 and SOD2, are both involved in oxidative metabolism, which is also known to be dysregulated in Rett Syndrome (De Felice et al., 2012). Thus, this CRISPR-enabled *Xenopus* model of Rett replicates many genetic and phenotypic features of the disease seen in humans.

### Computational discovery of drugs that reverse the network-level impact of Rett

In Rett, MeCP2 knockout results in widespread but subtle gene expression patterns, and most effects are not detectable using standard statistical analysis. Researchers have applied network-based analytical methods to uncover previously-hidden connections within RNA and protein interaction networks (Miller et al., 2019; Sanfeliu et al., 2019; Varderidou-Minasian et al., 2020), but these approaches have not been applied to Rett syndrome therapeutics discovery.

To leverage transcriptomics analysis for drug discovery, we developed a computational framework we termed nemoCAD (network model for causality-aware discovery)(**Fig. 2a**) that compares transcriptomic signatures of disease versus healthy control subjects to predict compounds from the Library of Integrated Network-Based Cellular Signatures (LINCS) database (“NIH LINCS Program,” n.d.), including FDA-approved drugs, which have a high likelihood to reverse the disease signature state back to a healthy state. Though signature-based tools have been previously developed to enable drug discovery (e.g., CMAP (Lamb et al., 2006), NicheNet (Browaeys et al., 2020), GPSnet (Cheng et al., 2019)), a major drawback is the impact of the expression of any single gene or protein on the resulting prediction. To be used effectively, these tools require well-matched disease datasets to the human cell line-based datasets present in LINCS, which may not be feasible for many diseases.

**Figure 2:**
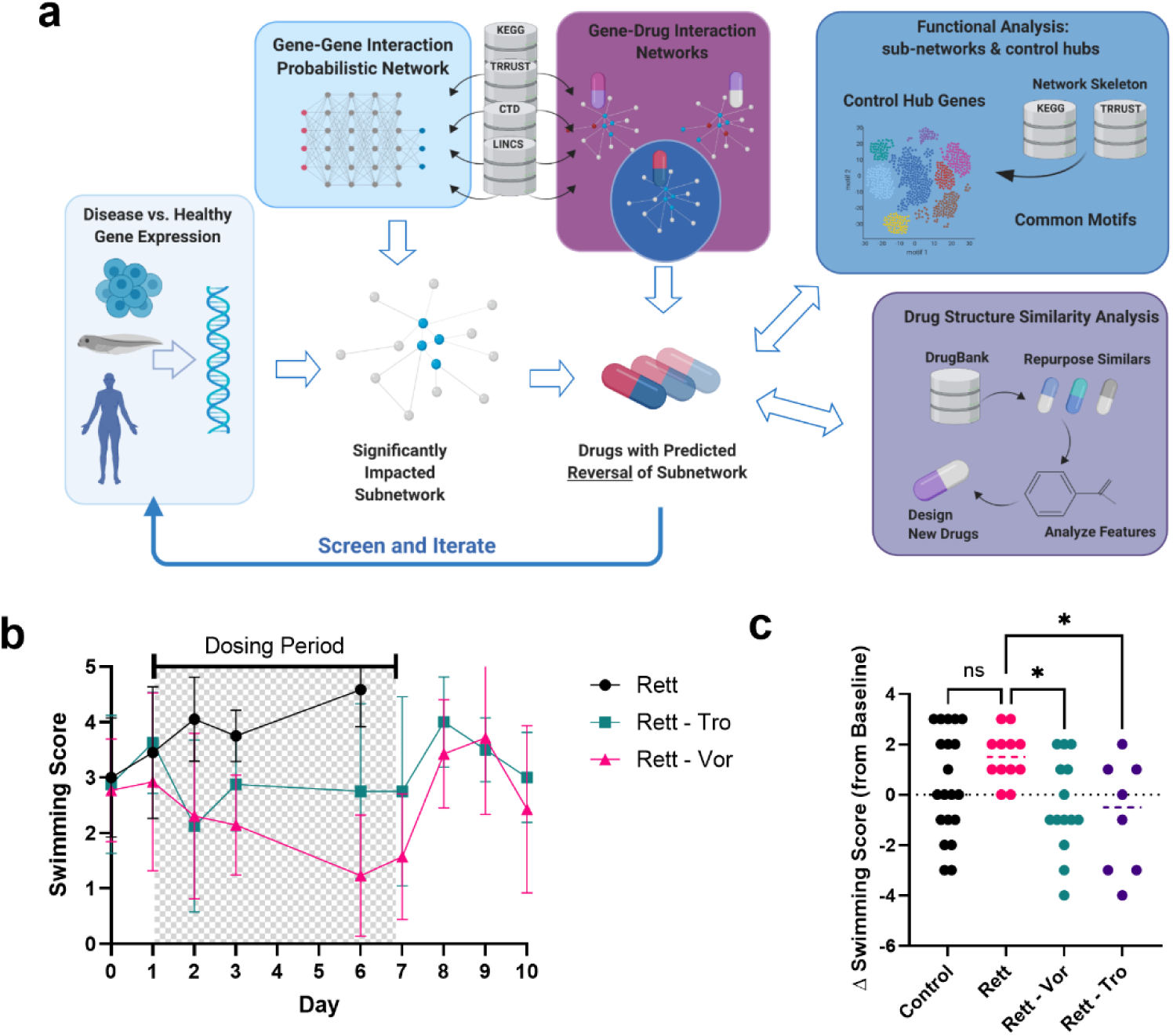
Network-based computational prediction of effective drugs to treat Rett syndrome in tadpole models. **a,** Network model for causality-aware discovery (nemoCAD) combines a directed gene-gene and drug-gene interaction network extracted from CTD, TRRUST, and KEGG databases with interaction probabilities inferred from single gene and drug perturbations in LINCS. Transcriptome data from any disease model or patient and corresponding control are used to identify the relevant subnetwork and disease-specific “node weights” that account for probabilities of up-/down-regulation of a gene. A drug-gene interaction probability matrix, inferred from LINCS, is computationally screened against the disease-specific subnetwork to identify compounds that significantly interact with the subnetwork and are ranked by their predicted ability to restore the disease transcriptome back to a healthy state based on single gene and gene network signatures. Downstream analyses can be performed on the resulting gene-gene interaction subnetwork by interrogating the underlying subnetwork structure to find control nodes and other network metrics. Additionally, the chemical structures of the predicted drugs can be clustered by structural similarity based on SMILES notation and annotated protein targets and pathways from DrugBank data. **b**, Graph showing relative effects on seizure score over 10 days of treatment in MeCP2 KD tadpoles of vehicle and 25 μM vorinostat (Rett - Vor) versus 70 μg/mL trofinetide (Rett - Tro), a clinical-stage drug with demonstrated efficacy (tadpoles per condition and timepoint: N = 12 Rett - Vor and Vehicle, N = 8 Rett - Tro; error bars indicate s.d.; ANOVA P = 0.028 Vehicle-treated MeCP2 KD tadpoles did not survive past day 3 of the treatment period in one study and past day 6 in a second study. **c**, Change in swimming score at day 6 of treatment vs. baseline (day 0) is significantly improved by both drugs. *, P < 0.05.

In contrast, we pursued an entirely different approach to transcriptomics signature analysis that is based on a Bayesian network encoding directed gene-gene and drug-gene interactions. By leveraging data in LINCS containing transcriptomics results following single gene knockout/knockdown and overexpression as well as addition of drug perturbagens in human cells, we calculated the probabilities of pairwise interactions among all of the genes and drugs in a human gene-gene and drug-gene interaction network whose structure was extracted from Comparative Toxicogenomics Database (CTD) (“The Comparative Toxicogenomics Database | CTD,” n.d.), KEGG (Kanehisa and Goto, 2000), and TRRUST (Han et al., 2018).

Unlike signature based computational tools that take gene expression signatures at face value, this probabilistic approach offers greater abstraction while prioritizing for the network impact of perturbations, including those caused by disease or addition of drugs, and thereby reduces the impact of tissue-specific or species-specific transcriptomic biases as well as the influence of the expression of any individual gene.

To identify drugs that might reverse disease states, we generated probabilistic network maps for each drug in the aggregate 19,800+ compound dataset. By inputting and comparing gene microarray data from MeCP2 knockdown tadpoles versus vehicle-injected controls, we identified sets of differentially expressed genes contained within the gene-gene interaction network with a p-value < 0.05 and a log2 fold change between 0.7 and 3.1 in 0.3-fold-change increments. For each fold-change, we generated a ranked list of compounds predicted to induce reversal of transcriptome-wide changes of: 1) single gene states using a cross-correlation score, 2) single gene states using a cross-entropy score, and 3) an overall gene network state score corresponding to the probability of involvement of the drug nodes in the identified subnetwork. These drug lists were combined to identify drugs that were ranked by being most consistently predicted to reverse the Rett transcriptome-wide changes and robust across the computational settings (**Supplementary Table 5**). Drugs were manually down-selected based on available toxicity data and to prioritize chemical diversity for initial screening. Rett tadpoles or controls were then dosed with lead compounds for 7 days via the culture medium beginning 1-2 days after the onset of symptoms and after collecting baseline swimming behavior data. Animal behaviors (swimming pattern abnormalities and seizure stages) were recorded every other day during drug exposure and washout phases, and the severity of seizure-like events was used as a screening metric.

These studies revealed that the FDA-approved drugs, vorinostat and ivermectin, demonstrated time-dependent reductions in seizure score that reversed after the drugs were washed out (**Fig. 2b**; **Supplementary Fig. 3**). However, we focused on vorinostat rather than ivermectin in all subsequent studies due to its higher predicted scoring consistency over time, more favorable reported pharmacokinetics, and its ability, albeit limited, to cross the blood-brain barrier. As a positive control with which to gauge relative efficacy, we also evaluated trofinetide, the synthetic IGF-1 C-terminal tripeptide. Although only moderately effective in recent Phase III Rett syndrome clinical trial top-line results that aligned with Phase II trial outcomes (Glaze et al., 2017), trofinetide represents the most relevant clinical-stage benchmark treatment. Importantly, in our Rett tadpole models, both trofinetide and vorinostat reduced seizure-like phenotypes and also increased viability (**Fig. 2b**). At day 6 of treatment, there was a statistically significant effect by both drugs on reducing swimming score relative to baseline before treatment onset (**Fig. 2c**).

As expected, no significant change was seen in WT and Rett vehicle-treated animals. This head-to-head comparison has supported the possibility that our *Xenopus* population-level model of Rett can offer predictive value for drug development.

To understand the protein targets of both drugs, we adapted thermal proteome profiling (Franken et al., 2015; Huber et al., 2015; Mateus et al., 2017) that identifies molecules that drugs bind directly to protein targets for use with a whole exothermic animal (*Xenopus* tadpole) as well as using isolated body segments (head, viscera, tail). While we were able to detect some of the known histone deacetylase (HDAC) targets of vorinostat, we also found that both this drug and ivermectin bind to multiple other proteins related more broadly to acetylation metabolism, beyond histones and HDACs (**Supplementary Table 6**). These findings suggest a potential mechanism of action for the Rett normalizing responses we observed that involves restoration of normal acetyl-CoA metabolism and post-translational acetylation.

As these observations initially aligned with results of prior studies of HDAC6 acetylation of tubulin in cells of Rett patients (Gold et al., 2015; Lebrun et al., 2021) and microtubule modulation in closely-related CDKL5 deficiency disorder (Barbiero et al., 2019), we analyzed tadpole tissues for α-tubulin acetylation and observed surprising bidirectional shifts in acetylation patterns depending on the tissue type. For example, α-tubulin was hypoacetylated in neurons in sections of the midbrain (**Supplementary Fig. 4a**) and hindbrain (**Fig. 3f**) and increased in GI tract (**Fig. 3g**) and olfactory multi-ciliated cells (**Supplementary Fig. 4b**) and following MeCP2 knockdown, resulting in more sparse but dense tangles. Higher resolution imaging revealed that the cilia were longer and misaligned on multiciliated cells compared to controls (**Fig. 3a**), and both length (**Fig. 3b**) and orientation (**Fig. 3c**) were restored by vorinostat treatment.

**Figure 3:**
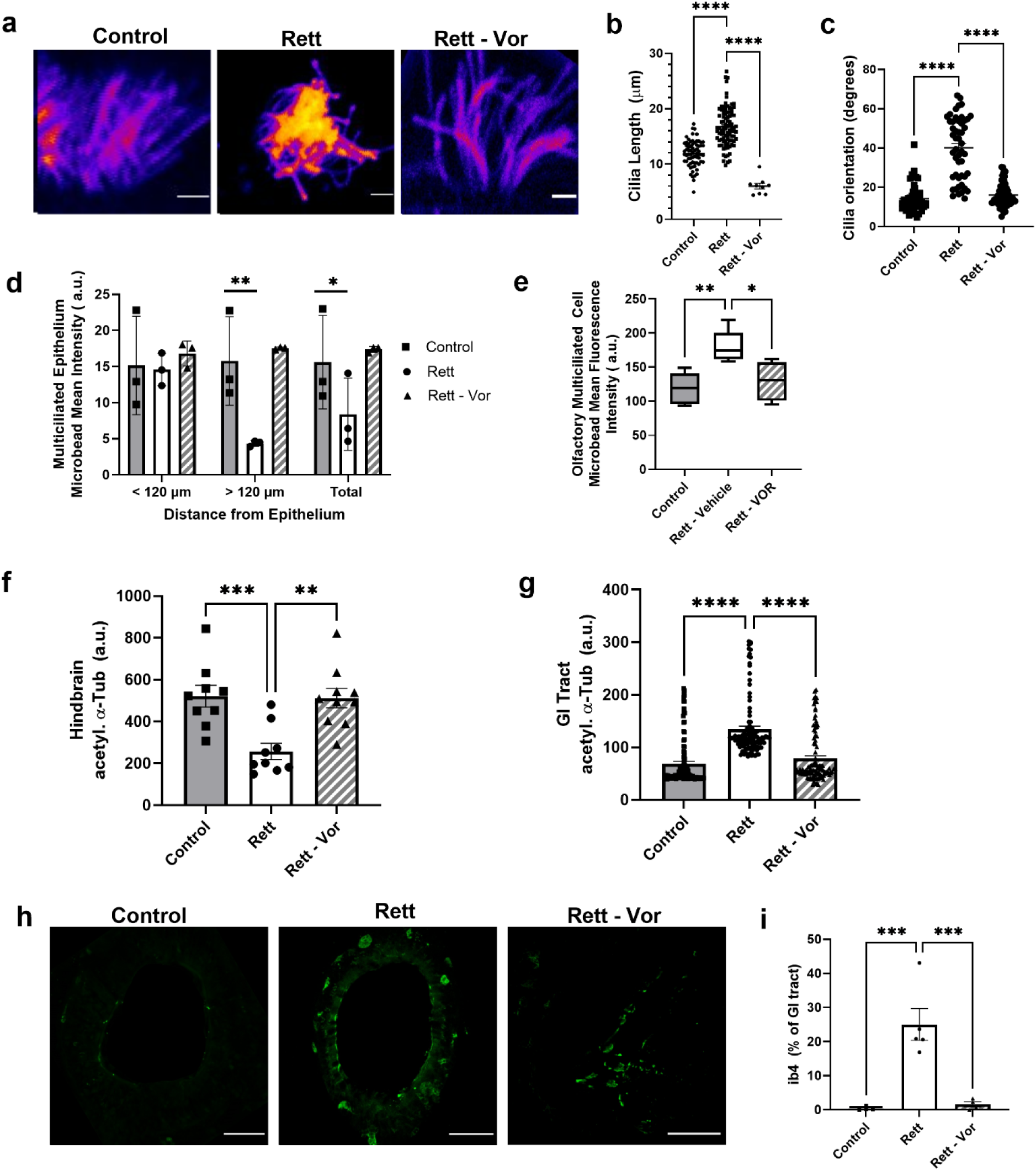
Vorinostat treatment of Rett tadpoles restores ciliary function, normalizes hypo- and hyperacetylated tubulin across tissues, and reduces GI tract inflammation. a, Olfactory cell ciliary abnormalities due to MeCP2 knockdown were restored by treatment with vorinostat (magenta-yellow colormap of fluorescence intensity, cilia stained for tubulin), significantly restoring their b, length and c, orientation (****, P < 0.0001). Functional rescue of cilia was also observed in both epithelial, d, and olfactory multiciliated cells in a microbead clearance assay (*, P < 0.05; **, P < 0.01). Despite the canonical HDACi activity of vorinostat, both hypoacetylated tubulin in the hindbrain, f, as well as hyperacetylated tubulin in the GI tract, g, was normalized. h, immunifluorescence images and i, plot showing that the % of ib4+ cells in the GI tract of tadpoles is increased in Rett tadpoles, which is indicative of inflammation and heightened pain response, and that this can be rescued by vorinostat treatment.

We then evaluated the impact of ciliary morphology and organization on ciliary function using fluorescent microparticles (**Supplemental Fig. 5**). The disruption of ciliary beating- induced flow on the surface of skin (**Fig. 3d**) and olfactory multiciliated cells (**Fig. 3e**) due to MeCP2 knockdown is reversed by vorinostat treatment, suggesting that ciliary function restoration is linked to ciliary function.

Due to the prevalence of GI symptoms in Rett patients and our observation of tubulin acetylation disruption and vorinostat-induced recovery in the GI tract (**Fig. 3g**), we further explored the role of MeCP2 knockdown in the gut. Staining for isolectin B4-positive (ib4+) cells to assess GI tract inflammation and nociceptive innervation, which is increased in Rett patients (Baikie et al., 2014; Downs et al., 2010), revealed that tadpoles exhibited significant ib4+ expression due to MeCP2 knockdown (**Fig. 3h,i)**, which vorinostat again reversed.

### Vorinostat normalizes the Rett phenotype in a mouse model

Because vorinostat was predicted to reverse the Rett state in our computational studies, we evaluated its efficacy in the MeCP2-deficient male (*MeCP2^-/Y^*) mouse Rett model that is commonly used in pre-clinical studies for Rett therapeutics, including trofinetide (Guy et al., 2001). We first treated younger animals starting at day 31 post-partum (p31) as was done in previous studies and confirmed that daily intraperitoneal (i.p.) administration of trofinetide (100 mg/kg) results in significant amelioration of multiple disease-related parameters, including diarrhea and motor function, measured by a reduction in the established Bird score (Guy et al., 2007) that incorporates a broad range of Rett-related CNS and non-CNS symptoms. When vorinostat (50 mg/kg) and trofinetide were similarly administered i.p. on a daily basis over two weeks and compared with vehicle control, we found that animals responded favorably to both drugs (**Fig. 4a**); however, treatment with vorinostat resulted in improved neurological function compared to trofinetide, as determined by measuring elevated plus maze (EPM) performance (**Fig. 4b**). Both drugs also resulted in enhanced spatial novelty Y-maze performance (**Fig. 4c**) and an improved diarrhea score (**Fig. 4d**).

**Figure 4:**
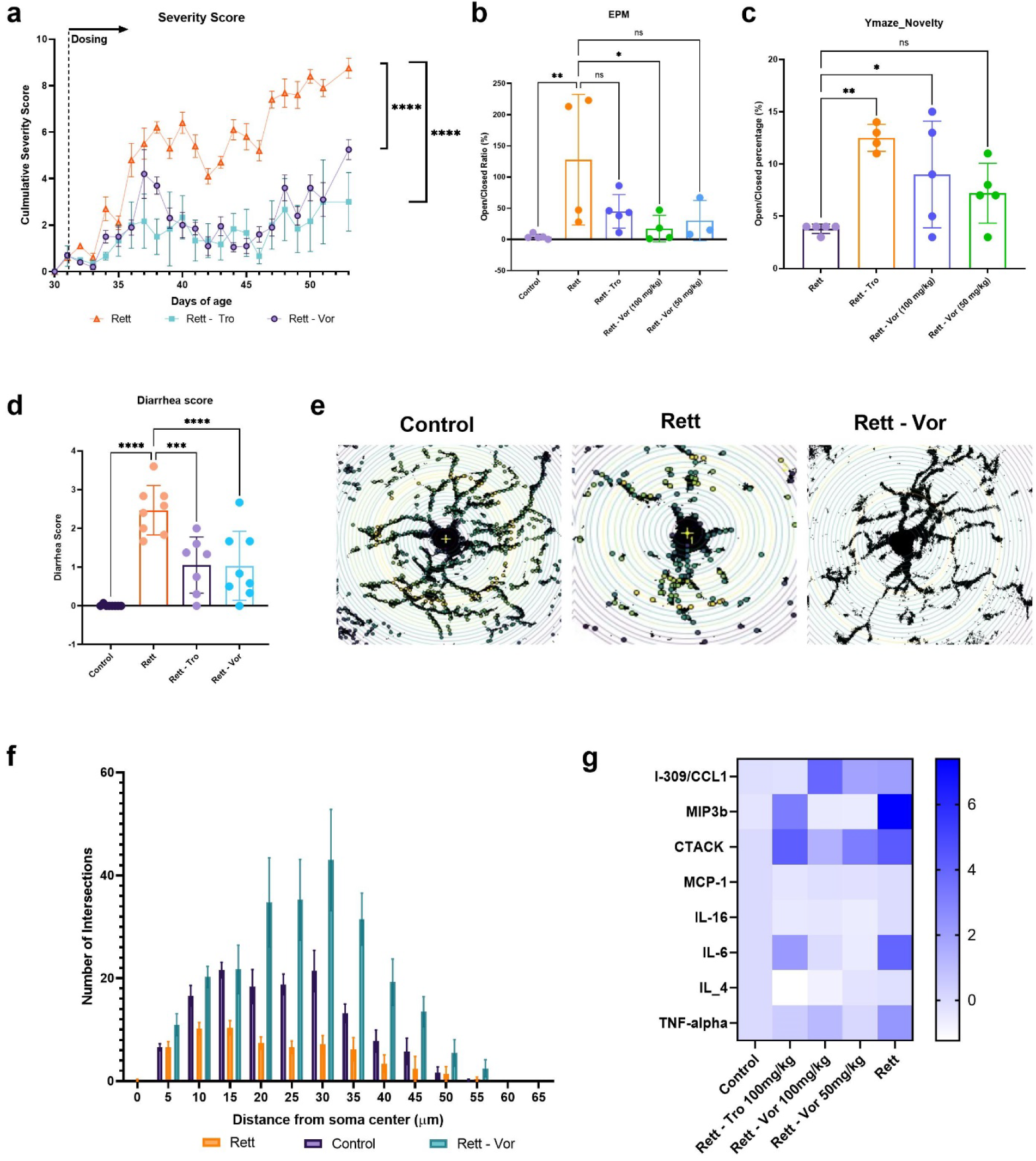
Vorinostat rescues multiple Rett syndrome-related CNS and somatic symptoms. **a,** Bird severity scores measured in MeCP2*^-/Y^* mice treated with vehicle (Rett), trofinetide (100 mg/kg), or vorinostat (50 mg/kg) from day 31 to 51 of age (ANOVA; ****, P < 0.0001). **b,** elevated-plus maze and, **c**, Y-novelty maze cognitive tests of MeCP2*^-/Y^* mice comparing vorinostat treatment efficacy to vehicle and trofinetide (ANOVA and Holm-Šídák test; *, P < 0.05; **, P < 0.01). **d**, diarrhea scored using a 0-3 scale (ANOVA and Holm-Šídák test; ***, P < 0.001; ****, P < 0.0001). Representative images of Sholl analysis of microglial arborization in mouse olfactory (**e**) and the Sholl analysis graph showing the branching profile of microglia (**f**) in control versus Rett mice treated with or without vorinostat (ANOVA and Holm-Šídák test; Control vs. Rett, P = 0.0004; Control vs. Vor, P < 0.0001). **g**, Levels of indicated cytokines measured in plasma of control mice versus Rett mice with or without treatment with trofinetide (Rett – Tro, 100 mg/kg), vorinostat (Rett – Vor, 50 mg/kg), or vehicle (Rett) for 3 weeks (mean values shown, N = 3, scale at right indicates z-score normalized to wild-type vehicle-treated littermates). The observed baseline inflammatory state in Rett mice agreed with published effects of MeCP2 knockdown(O’Driscoll et al., 2015).

As brain microglia have been implicated in Rett progression and therapeutic effects (Cronk et al., 2015; Derecki et al., 2012; Schafer et al., 2016) based on their role in neuroinflammation, waste removal, and synaptic pruning, we immuno-stained for microglia using ionized calcium binding adaptor molecule (Iba-1), a microglia-specific marker, in sections of MeCP2^-/y^ mouse brains. Using Sholl analysis to assess microglia morphology (Timmerman et al., 2018), we found that the MeCP2^-/y^ microglia exhibited significantly stunted morphology with fewer projection crossings compared to WT littermates, indicative of a heightened inflammatory state, which vorinostat restored (**Fig. 4e,f**). This result, together with the reduction of multiple serum inflammatory markers, including TNF-α, IL-4, IL-6, IL-16, CCL2, MIP3b, and CCl1 (**Fig. 4g**), suggests that animals with Rett experience a heightened inflammatory state, and that treatment with vorinostat can suppress this inflammation.

### Oral vorinostat exhibits broad efficacy in Rett mice even after onset of symptoms

Pre-clinical studies for Rett therapies routinely dose MeCP2^-/y^ mice prior to the onset of symptoms (Castro et al., 2014; Szczesna et al., 2014) due to the rapid onset of symptoms and short lifespan of animals, in contrast to patients who seek clinical care due to developmental deficits which may not be diagnosed for several years before treatment could be initiated (Tarquinio et al., 2015). To better mimic this more relevant clinical scenario, male MeCP2^-/y^ mice were dosed approximately one week after the onset of symptoms. Also, because long term treatment of Rett patients would be best served by oral administration, and due to the fact that our results showed a reduction in diarrhea in mice treated with vorinostat via daily i.p., we tested the efficacy of an oral formulation of vorinostat. Interestingly, when we administered trofinetide i.p. after symptoms developed in this model, it was found to be ineffective based on overall Rett score, similar to the moderate efficacy observed in human clinical trials, whereas oral vorinostat (50 mg/kg) prevented significant worsening of the symptom severity score (**Fig. 5a**), ameliorated weight gain, and increased performance in EPM (**Fig. 5b**) and Y mazes (**Fig. 5c**). Animals treated with oral vorinostat showed complete survival after 3 weeks of treatment, which is comparable to what would be expected for vehicle-treated Rett animals for a study of this duration, whereas only ∼ 60% of trofinetide-treated Rett animals survived (**Fig. 5d**). To our knowledge, this represents the first effective treatment of MeCP2^-/y^ mice when treatment is initiated *after* the onset of symptoms.

**Figure 5:**
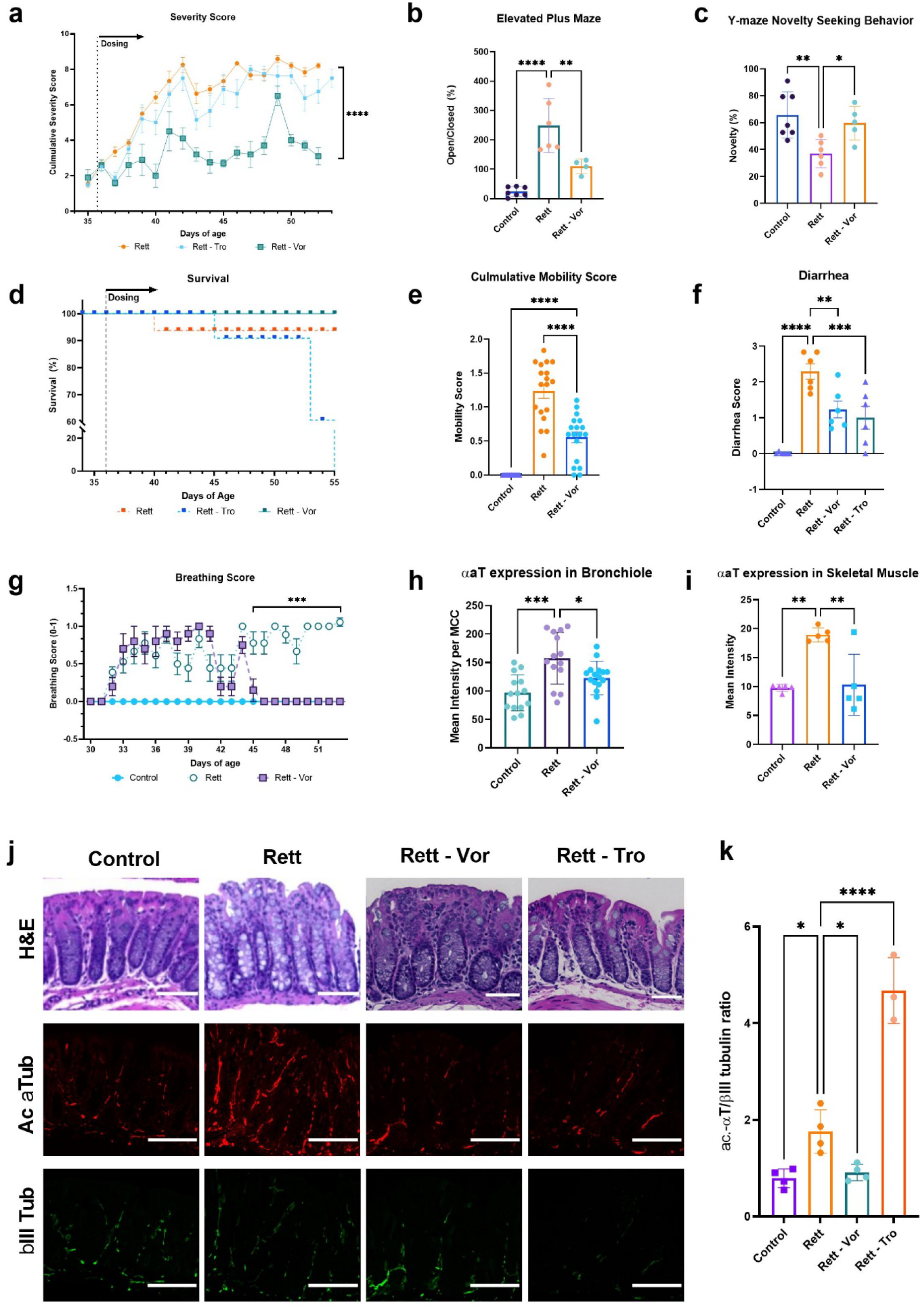
Oral administration of vorinostat in MeCP2^-/y^ mice after the onset of Rett symptoms rescues CNS and somatic symptoms. **a,** Bird severity scores measured in MeCP2*^-/Y^* mice treated with vehicle (Rett), trofinetide (100 mg/kg i.p.), or oral vorinostat (50 mg/kg) from day 36, after onset of symptoms, showing significant suppression of symptoms by vorinostat treatment while trofinetide’s effects were indistinguishable from the vehicle control (ANOVA, ****, P < 0.0001). **b,** elevated-plus maze and, **c**, Y-novelty maze cognitive tests of MeCP2*^-/Y^* mice comparing vorinostat treatment efficacy to vehicle and trofinetide (ANOVA and Dunnett’s multiple comparison test; *, P < 0.05; **, P < 0.01; ****, P < 0.0001). **d**, survival curve of animals in this study. **e**, mobility scored using a 0-2 scale and **f**, diarrhea scored using a 0-3 scale (ANOVA and Holm-Šídák test; **, P < 0.01 ***, P < 0.001; ****, P < 0.0001). **g**, breathing difficulty evaluated using a 0-1 score (ANOVA and Dunnett’s multiple comparison test; ***, P < 0.001). Graphs showing levels of hyperacetylated α-tubulin in cells of the multi-ciliated cells in the bronchiolar epithelium of the lung (**h**) and skeletal muscle (**i**) in mice treated as described in **a** (data are presented as mean ± s.d.; *, P < 0.05; **, P < 0.01; ***, P < 0.001). **j**, Hematoxylin and eosin (H&E) stained histological sections of colon from the control and drug-treated mice (top row) and immunofluorescent staining for acetylated α-tubulin (middle) and βIII-tubulin (bottom) in these sections. **k**, Graph showing changes in the βIII-tubulin ratio in drug-treated versus control colon tissues (*, P < 0.05; ****, P < 0.0001).

Rett patients and mouse Rett models exhibit significant declines in fine motor and gait control as well as GI tract dysfunction(Hagberg, 2002; Motil et al., 2012). Importantly, the deficiencies in mobility, gait, and hindlimb clasping, as well as increased diarrhea we also observed in MeCP2^-/y^ animals were again rescued by vorinostat treatment (**Fig. 5e,f**). Breathing also was significantly improved following vorinostat treatment compared to trofinetide and vehicle treatments (**Fig. 5g**). Given the putative effects of vorinostat on acetylation metabolism and post-translational modification of tubulin and its disruption of ciliary function identified in tadpoles, we also analyzed α-tubulin acetylation in the multi-ciliated cells in mouse lung bronchioles, which revealed an overall increase of hyperacetylated α-tubulin in MeCP2^-/y^ animals that vorinostat again restored (**Fig. 5h**). In addition, we explored the potential link of tubulin acetylation with muscle function and found that vorinostat treatment can rescue the disrupted α-tubulin acetylation observed in femoral muscle sections (**Fig. 5i**). Increased acetylation due to the loss of MeCP2 may play a role in muscle function as hyperacetylation has been shown to increase stiffness and resistance in striatal muscles *in vitro* (Coleman et al., 2021).

We continued to evaluate the possible impact of tubulin acetylation by examining the colon, a biomechanically-active organ that is often distended in Rett patients and may play a major role in digestive tract disruption that is a major complaint of these patients clinically (Motil et al., 2012). Initial histological analysis of the colon indicated a greater degree of vacuolization and heightened neutrophil infiltrate in MeCP2-null animals, which vorinostat and trofinetide reversed (**Fig. 5j**). Staining of these sections for βIII- and acetylated α-tubulin also revealed that acetylated α-tubulin is significantly more colocalized with neuronal staining in the colon in MeCP2-null mice, and that this can be normalized by treatment with either vorinostat or trofinetide (**Fig. 5j**). However, while vorinostat also normalized the ratio of acetylated α-tubulin to βIII-tubulin, it was unexpectedly worsened by trofinetide (**Fig. 5j,k**). While the developing GI tract of tadpoles prevented detailed analysis of their enteric neurons, the tubulin acetylation- normalizing effect of vorinostat identified by network analysis in *Xenopus* clearly translated to this mammalian (mouse) model at the cellular level.

These data show that oral dosing of vorinostat improved classical disease outcome metrics in MeCP2-null male mice and it ameliorated microglial dysfunction and hyperacetylation of tubulin in their GI and respiratory tracts as well as in skeletal muscle. Vorinostat also was recently shown to be effective in treating Fragile X syndrome (Ding et al., 2021), which is closely related to Rett, sharing both X-linked mutations and symptoms of autism, although it is caused by a mutation in the FMR1 gene. Interestingly, we identified a loss of gene network connectivity with FMR1 in the Rett-related *Xenopus* gene network (**Fig. 1h**), and genes with the maximum variation and fold change differences in humans with Fragile X also occurred in pathway classes involved in oxidative stress and multiple biosynthesis pathways that align with the major regulators we identified by the cross-species MeCP2 mutant network analysis (**Supplementary Figure 6a** and **Supplementary Tables 7 and 8**). Moreover, when we used nemoCAD to predict drugs that might reverse the Fragile X genotype in humans, both the compound classes and gene targets were similar to those predicted for *Xenopus* with MeCP2 knockdown, including a high number of PPAR receptor agonists, HDAC inhibitors, and cyclooxygenase inhibitors (**Supplementary Figure 6b,c**). Thus, these computational results suggest more broadly that developmental therapeutics approaches for neurodevelopmental disorders with similar broad transcriptional effects may benefit from using this form of gene interaction network analysis to identify network-level signatures characteristic of specific diseases and to predict drugs that may reverse those changes.

## DISCUSSION

Taken together, our results demonstrate the value of combining computational network analysis-based drug repurposing algorithms with CRISPR technology to rapidly build novel *in vivo* disease models using *X. laevis* that can be used to generate data and carry out screening of drugs with activities of interest predicted computationally. The *Xenopus* tadpole model developed here, which was generated in our laboratory within 3 weeks, exhibits a large degree of genetic and phenotypic heterogeneity that reduces risks associated with computational and screening outcomes by better capturing the disease state. Importantly, the *Xenopus* results translated well to a mouse Rett model, and hence the tadpole model offers a faster, lower-cost, and potentially much higher throughput alternative to mammalian models. In addition, our target-agnostic computational approach enabled rapid discovery of novel mechanisms, including the counterintuitive ability of an HDAC inhibitor to normalize both hypoacetylated tubulin in the CNS and hyperacetylated tubulin in other organs, as well as identify drugs that have broad efficacy across multiple organs. This approach led to the first demonstration of efficacy in the mouse MeCP2 knockout model of Rett syndrome even when treated after the onset of symptoms, in contrast with the lack of efficacy of trofinetide. Given the modest efficacy of trofinetide in the clinic, conducting pre-clinical studies to restore health after the onset of symptoms as in our *Xenopus* screen may be a more robust benchmark for efficacy in complex CNS disorders. The drug discovery platform presented here offers Rett and other neurodevelopmental patients the possibility of single small molecule therapeutics that address multiple fundamental disease-modifying pathways with a broad clinical impact. Furthermore, by identifying and rapidly evaluating existing drugs, potentially even in patients, our platform enables the discovery of hidden therapeutic mechanisms for new drug development.

## ACKNOWLEDGEMENTS

We would like to thank Dr. Barbara J. Caldarone for guidance and assistance with the mouse neurobehavioral assays, Dr. Amanda Graveline, Sarai Bardales, Andyna Vernet and Melissa Sanchez-Ventura for their assistance with mouse studies, Thomas C. Ferrante for guidance and assistance with image acquisition and analysis, Dr. Lay-Hong Ang and Suzanne White for tissue immunostaining support, Dr. Aurelien Begue and Mahmoud El-Rifai for their assistance with RNAscope experiments, and Erin Switzer and Emma Lederer for help with *Xenopus* embryo fertilization and husbandry. All behavioral experiments were conducted at the HMS Mouse Behavioral Core facility. M.L. gratefully acknowledges support of the Elisabeth Giauque Trust, London. We are grateful for funding from the Wyss Institute for Biologically Inspired Engineering at Harvard University through Validation Project support.

## AUTHOR CONTRIBUTIONS

RN, FV, and DEI conceived of and directed the work. RN, TL, MS, FV, EG, KS, KV, and AV conducted animal studies and in vitro analyses. FV, MS, and VK developed and performed CRISPR generation of Rett tadpole models with guidance by ML. SK, MS, SL, VC, and TT performed bioinformatic analysis and computational drug prediction. SL and RN designed and developed nemoCAD software. JRT guided GI tract and lung immunostaining studies and evaluated the histology and tissue staining. RN and DEI wrote the manuscript. All authors reviewed and provided input for the manuscript.

## DATA AVAILABILITY

All data used in this study, including the *Xenopus* microarray data, are publicly available using accession codes listed in **Supplementary Table 2**.

## CODE AVAILABILITY

Code is available upon reasonable request to the corresponding author.

