## Supplemental Information for "Target-agnostic discovery of Rett Syndrome therapeutics by coupling computational network analysis and CRISPR-enabled *in vivo* disease modeling"

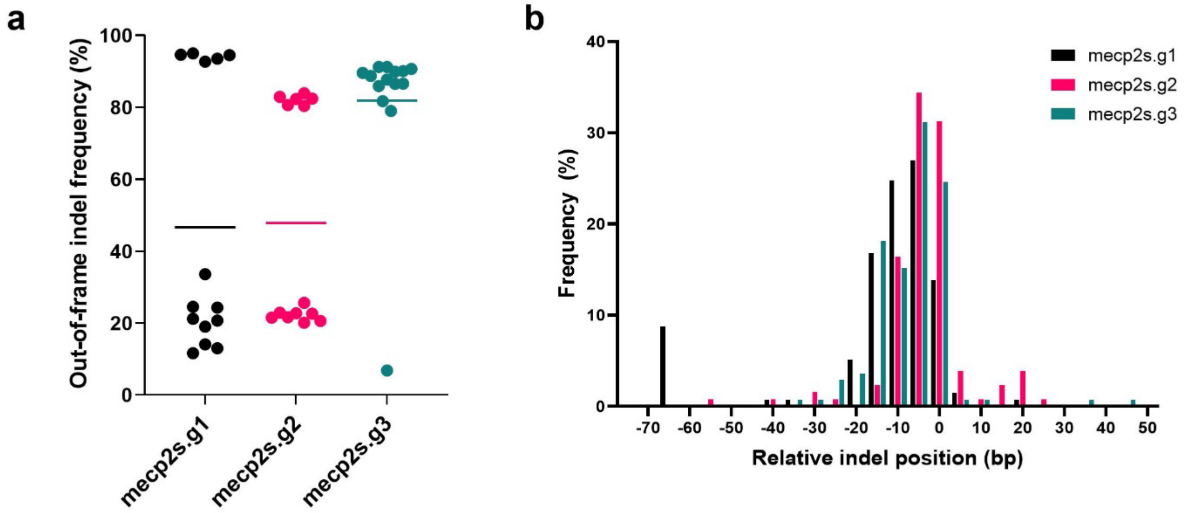


**Supplementary Figure 1: CRISPR-editing the mecp2 gene in Xenopus following fertilization generates a heterogenous population of MeCP2-deficient tadpoles.**

**a**, Frequency of indels determined using IDAA analysis from a subset of 14 tadpoles at 3 target sites on the MeCP2.S gene demonstrating the mosaicism and broad range of editing efficiency driving diverse phenotype severity. **b**. IDAA analysis further demonstrating the variability in indel position due to the stochastic nature of post-fertilization CRISPR editing that drives observed phenotypic variability.

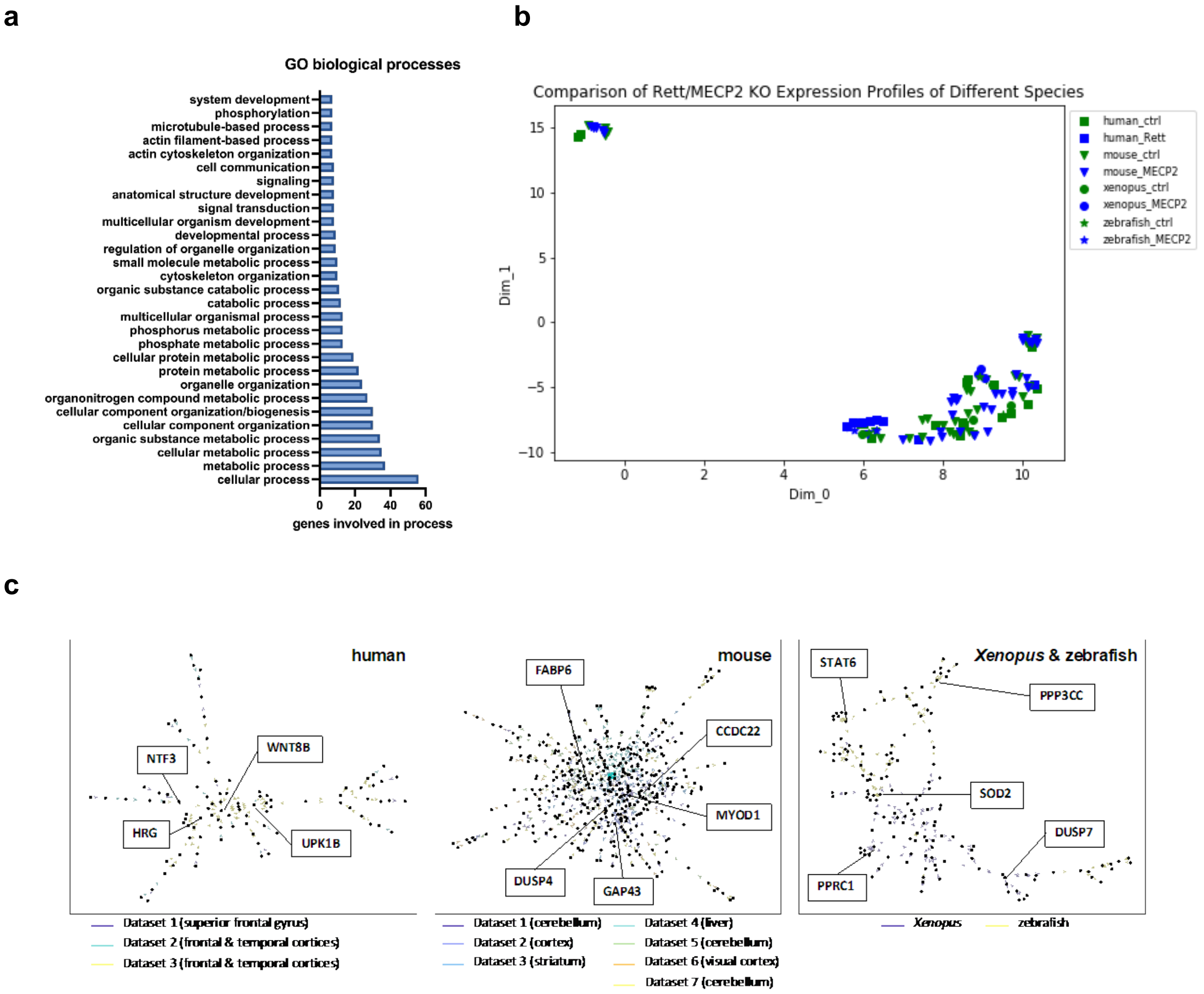


**Supplementary Figure 2: Comparative analysis of animal models of Rett syndrome and human patients.**

**a**, Gene ontology (GO) biological processes identified for the 70 statistically-significant genes in MeCP2 knockdown tadpoles compared to controls. **b**, UMAP clustering of gene expression signatures comparing the Xenopus model from this work to publicly available datasets showing Rett syndrome clinical or animal model data (blue) compared to controls (green). **c**, network analysis identified human, mouse, and whole-organism (Xenopus tapdole, zebrafish) unique network structures.

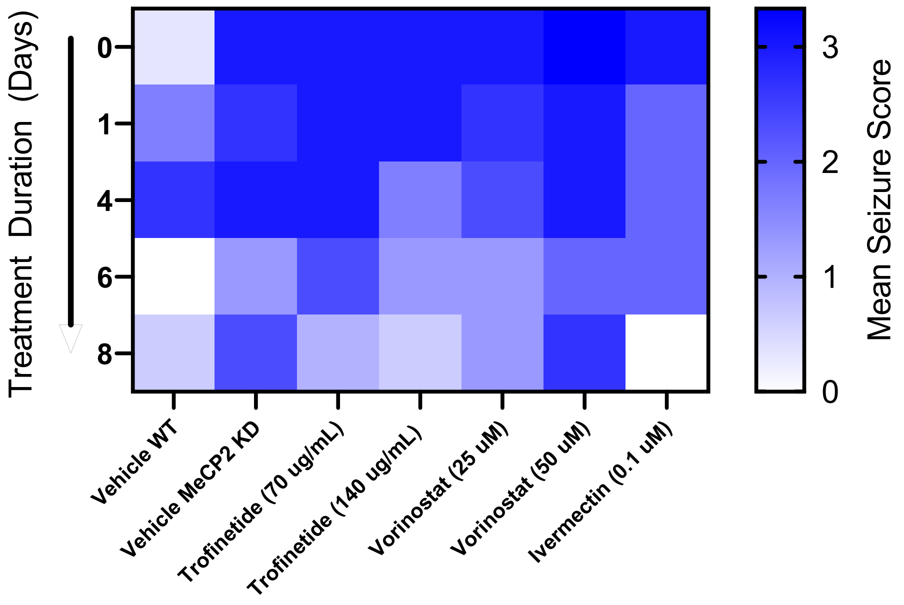


**Figure 3: *Xenopus laevis* Rett syndrome screening results.** Seizure scores for wildtype tadpoles with vehicle alone (WT), MeCP2 knockdown tadpoles (MeCP2 KD), and MeCP2 KD animals treated with indicated doses of trofinetide (Tro), vorinostat (Vor), or ivermectin (Ive) for 0 to 8 days duration starting one week post-fertilization (color scale bar at right indicates mean seizure score).

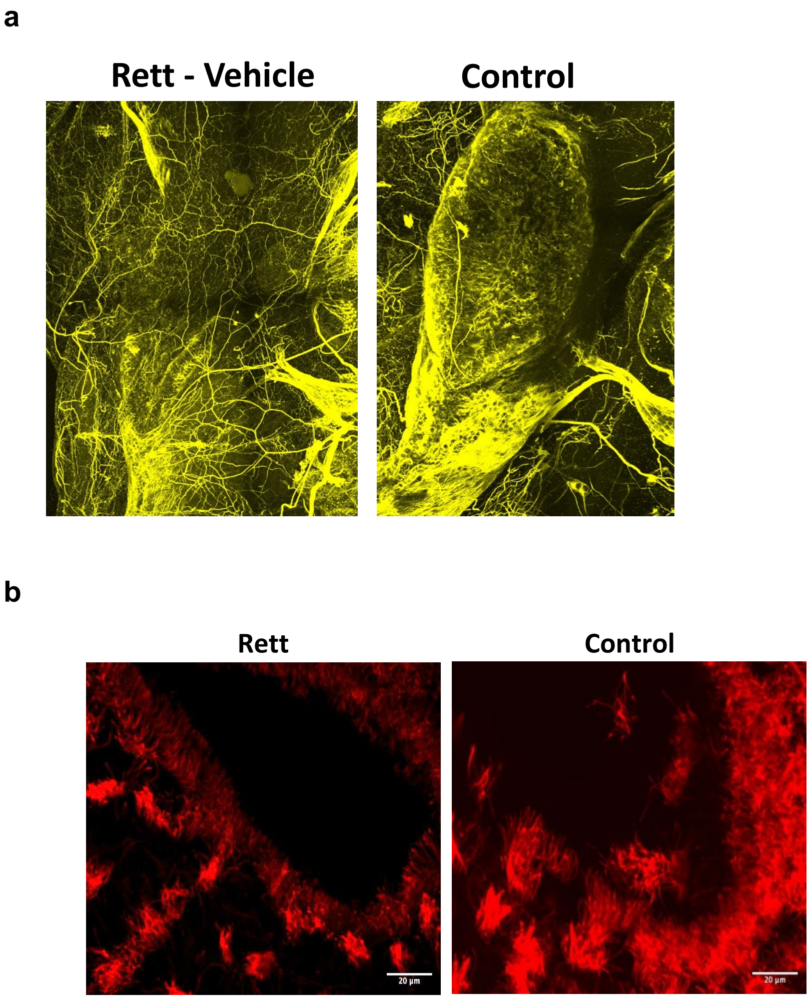


**Supplementary Figure 4: The bidirectional impact of MeCP2 demonstrated in decreased acetylation of α-tubulin in the brain of the tadpole model of Rett syndrome and increased acetylation in mucociliary multi-ciliated cells.**

Acetylation of neuronal α-tubulin is reduced in Rett tadpole midbrain, **a**, and increased in olfactory multi-ciliated cells, **b**, resulting in more compact, tangled cilia. Immunostaining against acetylated α-tubulin in both images. Scale bars 20 µm.

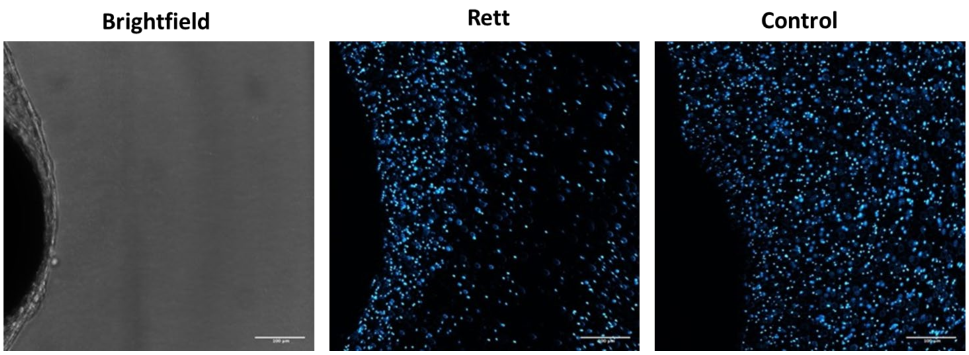


**Supplementary Figure 5: Ciliary function in tadpole Rett model is rescued by vorinostat treatment.**

To visualize cilia-driven epidermal flow, fluorescent microbeads were measured using fluorescence imaging at steady state relative to the proximity of tadpole epidermal multi-ciliated cells (MCC) over a period of 5 minutes. Scale bars 200 μm.

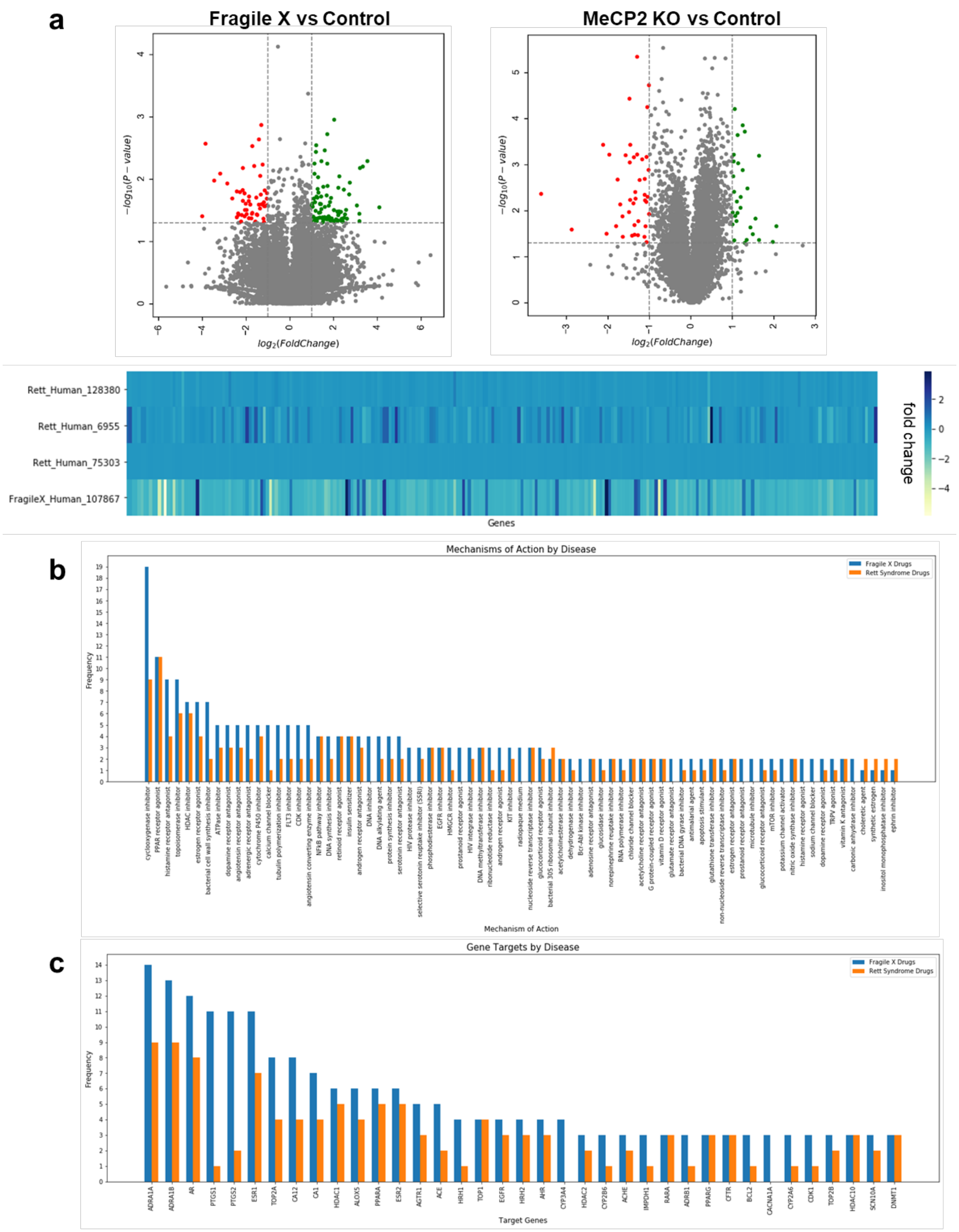

a
Fragile X vs Control
MeCP2 KO vs Control
$-log_{10}(P-value)$
$log_2(FoldChange)$
Rett_Human_128380
Rett_Human_6955
Rett_Human_75303
FragileX_Human_107867
Genes
fold change
b
Mechanisms of Action by Disease
Fragile X Drugs
Rett Syndrome Drugs
Frequency
Mechanism of Action
cyclooxygenase inhibitor
PPAR receptor agonist
histamine receptor antagonist
topoisomerase inhibitor
HDAC inhibitor
estrogen receptor agonist
bacterial cell wall synthesis inhibitor
ATPase inhibitor
dopamine receptor antagonist
angiotensin receptor antagonist
adrenergic receptor antagonist
cytochrome P450 inhibitor
calcium channel blocker
tubulin polymerization inhibitor
FLT3 inhibitor
CDK inhibitor
angiotensin converting enzyme inhibitor
NFkB pathway inhibitor
DNA synthesis inhibitor
retinoid receptor agonist
insulin sensitizer
androgen receptor antagonist
DNA inhibitor
DNA alkylating agent
protein synthesis inhibitor
serotonin receptor antagonist
HIV protease inhibitor
selective serotonin reuptake inhibitor (SSRI)
phosphodiesterase inhibitor
EGFR inhibitor
HMGCR inhibitor
prostanoid receptor agonist
HIV integrase inhibitor
DNA methyltransferase inhibitor
ribonucleotide reductase inhibitor
androgen receptor agonist
KIT inhibitor
radiopaque medium
nucleoside reverse transcriptase inhibitor
glucocorticoid receptor agonist
bacterial 30S ribosomal subunit inhibitor
acetylcholinesterase inhibitor
dehydrogenase inhibitor
Bcr-Abl kinase inhibitor
adenosine receptor antagonist
glucosidase inhibitor
norepinephrine reuptake inhibitor
RNA polymerase inhibitor
chloride channel blocker
acetylcholine receptor antagonist
G protein-coupled receptor agonist
vitamin D receptor agonist
glutamate receptor antagonist
bacterial DNA gyrase inhibitor
antimalarial agent
apoptosis stimulant
glutathione transferase inhibitor
non-nucleoside reverse transcriptase inhibitor
estrogen receptor antagonist
prostanoid receptor antagonist
microtubule inhibitor
glucocorticoid receptor antagonist
mTOR inhibitor
potassium channel activator
nitric oxide synthase inhibitor
histamine receptor agonist
sodium channel blocker
dopamine receptor agonist
TRPV agonist
vitamin K antagonist
carbonic anhydrase inhibitor
choleretic agent
synthetic estrogen
inositol monophosphatase inhibitor
ephrin inhibitor
c
Gene Targets by Disease
Fragile X Drugs
Rett Syndrome Drugs
Frequency
Target Genes
ADRA1A
ADRA1B
AR
PTGS1
PTGS2
ESR1
TOP2A
CA12
CA1
HDAC1
ALOX5
PPARA
ESR2
AGTR1
ACE
HRH1
TOP1
EGFR
HRH2
AHR
CYP3A4
HDAC2
CYP2B6
ACHE
IMPDH1
RARA
ADRB1
PPARG
CFTR
BCL2
CACNA1A
CYP2A6
CDK1
TOP2B
HDAC10
SCN10A
DNMT1

**Supplementary Figure 6: Comparison of Fragile X syndrome transcriptome data to Rett syndrome reveals many similarities in gene expression and predicted drug mechanisms of action.**

Volcano plots of Fragile X and Rett syndrome gene expression reveal similar low numbers of significantly impacted genes that are overall reduced in magnitude for Fragile X relative to Rett syndrome, **a**. Drugs predicted using nemoCAD to restore Fragile X or Rett syndrome gene expression signatures to a healthy state share many mechanisms of action, **b**, and target genes, **c**.

### Supplementary Table 1. Genes that undergo significant change following MeCP2 knockdown compared to control

| Gene | log2 Fold Change | Adjusted p-value |
|---|---|---|
| GLIS2 | -0.34078 | 0.01077909 |
| SUCLA2 | 0.66681838 | 0.01077909 |
| FAM213B | 1.29421399 | 0.01077909 |
| GDPD1 | -0.5772415 | 0.01077909 |
| EPS8 | -0.8369083 | 0.01077909 |
| SLC9A3R1 | -0.5171517 | 0.01470034 |
| HSPB1 | 0.70090785 | 0.02148022 |
| UBQLN4 | -0.290626 | 0.02561644 |
| KCTD2 | -0.4091327 | 0.02561644 |
| TCTN1 | -0.6492749 | 0.02561644 |
| NUP210 | 0.57992661 | 0.02561644 |
| SLC25A32 | 0.76237415 | 0.02561644 |
| MTR | 1.00955786 | 0.02561644 |
| NDUFS7 | 0.22687485 | 0.0267425 |
| MYSM1 | -0.3715985 | 0.0267425 |
| ADCK1 | 1.47869428 | 0.0267425 |
| TIGAR | 0.66837166 | 0.02858852 |
| MOB2 | -0.4605303 | 0.02987123 |
| LONRF1 | 0.86049786 | 0.02987123 |
| ABHD17B | -0.2353246 | 0.02987123 |
| SLC37A2 | -1.0614915 | 0.02987123 |
| CLN6 | -0.3286455 | 0.02987123 |
| SLC35A4 | 1.04945715 | 0.02987123 |
| HMG20A | -0.4609388 | 0.03171146 |
| ETFDH | 0.52905831 | 0.03171146 |
| MPRIP | -0.4534216 | 0.03171146 |
| SSH3 | -0.3707231 | 0.03171146 |
| GOLGA5 | -0.2633019 | 0.03171146 |
| NTAN1 | 0.95594894 | 0.03171146 |
| NFKBIE | -0.4704496 | 0.03385416 |
| AKT1S1 | -0.7300014 | 0.03659645 |
| METTL23 | -0.7758509 | 0.04317334 |
| FAM921 | -0.3385863 | 0.04365897 |
| CFAP299 | -1.2521905 | 0.04365897 |
| SLC25A40 | 0.78718125 | 0.04365897 |
| EFS | -0.8743784 | 0.04464713 |
| ACTR3 | -0.1984858 | 0.04465646 |
| GLRX | 0.46728133 | 0.04465646 |
| TRIM72 | -0.6729334 | 0.04465646 |
| NFS1 | 0.63952963 | 0.04465646 |
| MOGAT1 | -0.4865663 | 0.04465646 |
| DDHD2 | -0.6861136 | 0.04465646 |
| ZNRD1 | -0.4538309 | 0.04465646 |
| SCRN2 | -0.8905708 | 0.04465646 |
| GGH | 0.57543546 | 0.04465646 |
| CLK2 | -0.7422218 | 0.04465646 |
| CCDC160 | -1.3031789 | 0.04465646 |
| FTSJ1 | -0.6301232 | 0.04465646 |
| CEP68 | -0.4567722 | 0.04465646 |
| IFT22 | -0.9132039 | 0.04465646 |
| TARBP1 | 0.86779331 | 0.04465646 |
| LOC100145070 | 0.13939757 | 0.04465646 |
| MOSPD1 | -0.4520379 | 0.04550288 |
| ABTB1 | -0.5997957 | 0.04550288 |
| CYP26A1 | -1.1296027 | 0.04550288 |
| PAOX | -0.4907751 | 0.04550288 |
| UQCRC1 | 0.22950474 | 0.0456716 |
| HEBP2 | 0.88530432 | 0.0456716 |
| SLC25A46 | -0.4125815 | 0.0456716 |
| LOC100496109 | 0.69713913 | 0.0456716 |
| ZMYND10 | 0.60297138 | 0.0456716 |
| HNF4B | -0.7665617 | 0.04623931 |
| KANSL1 | -0.4998616 | 0.04623931 |
| SLC41A1 | 0.83533564 | 0.04623931 |
| ABI1 | -0.8988648 | 0.04890499 |
| THOC3 | -0.2528048 | 0.04890499 |
| HUS1 | -0.3286943 | 0.04890499 |
| NEK6 | -0.4961035 | 0.04890499 |
| TMEM40 | -0.3758727 | 0.04890499 |
| ZFP64 | -0.7143661 | 0.04890499 |

**Supplementary Table 2. Datasets used in cross-species transcriptomics analysis**

| dataset identifier | species | patient or model details | tissue | reference |
|---|---|---|---|---|
| GSE6955 | human | Rett syndrome (females aged 2-10y) | brain (superior frontal gyrus) | Deng et al. 2007 |
| E-GEOD-75303 | human | Rett syndrome (females aged 17-21y) | brain (frontal & temporal cortices) | unpublished |
| GSE128380 | human | Rett syndrome (females aged 16-32y) | brain (frontal & temporal cortices) | Aldinger et al. 2020 |
| E-GEOD-42895 | mouse | MeCP2 KO (males aged 8 weeks) | brain & liver (brain: striatum) | Zhao et al. 2013 |
| GSE67294 | mouse | MeCP2 KO (males aged 8-9 weeks) | brain (visual cortex) | Gabel et al. 2015 |
| GSE96684 | mouse | MeCP2 KO (males, postnatal day 60) | brain (whole cortex) | Pacheco et al. 2017 |
| GSE105045 | mouse | MeCP2 KO (males aged 8-9 weeks) | brain (cerebellum) | Raman et al. 2018 |
| GSE112663 | mouse | conditional MECP2 KO mice (males aged 22 weeks) | brain (cerebellum & cerebral cortex) | unpublished |
| GSE199049 | Xenopus laevis | MeCP2 KD (Stage 50 tadpoles) | whole organism | current work |
| GSE80348 | zebrafish | MeCP2 KO (6 hours post-fertilization) | whole organism | van der Vaart et al. 2017 |

**Supplementary Table 3: Cross-species network hubs**

| Species | Hub Genes | Primary Function | In-degree | Out-degree |
|---|---|---|---|---|
| human | WNT8B | brain development | 17 | 1 |
| | UPK1B | signal transduction | 15 | 0 |
| | HRG | blood clotting | 8 | 0 |
| | SERPIND1 | inflammation & clotting | 5 | 2 |
| | NTF3 | nerve maintenance | 4 | 1 |
| | GNPAT | metabolism & biosynthesis | 0 | 4 |
| | FZD7 | Wnt signaling | 2 | 4 |
| | PAX8 | embryogenesis & cancer | 0 | 3 |
| | MRPS31 | protein synthesis | 0 | 3 |
| | GCLM | glutamate-cysteine metabolism | 0 | 3 |
| mouse | DUSP4 | signal transduction | 11 | 4 |
| | GAP43 | nerve growth | 10 | 0 |
| | MYOD1 | muscle cell differentiation | 9 | 1 |
| | FABP6 | fatty acid metabolism | 9 | 2 |
| | CCDC22 | calcium signaling | 9 | 3 |
| | FDPS | isoprenoid biosynthesis | 3 | 8 |
| | CH25H | lipid metabolism | 3 | 8 |
| | SLBP | histone mRNA processing | 0 | 7 |
| | RGS4 | signal transduction | 2 | 7 |

| | | | | |
|---|---|---|---|---|
| | NR4A3 | transcriptional activation | 2 | 7 |
| | HGS | signal transduction | 5 | 0 |
| | MED8 | transcriptional activation | 4 | 0 |
| | GLDC | protein degradation | 4 | 0 |
| | FGF9 | nervous system development | 4 | 1 |
| *Xenopus* & zebrafish | ELL2 | immunoglobulin processing | 4 | 0 |
| | SOD2 | oxidative stress | 3 | 10 |
| | STAT6 | signal transduction | 0 | 8 |
| | PPRC1 | mitochondrial biogenesis | 0 | 8 |
| | TBC1D4 | glucose homeostasis | 0 | 7 |
| | PPP3CC | calcium signaling | 0 | 7 |
| | WNT8B | brain development | 20 | 2 |
| | UPK1B | signal transduction | 15 | 5 |
| | MYOD1 | muscle cell differentiation | 12 | 1 |
| | GAP43 | nerve growth | 11 | 0 |
| across species | DUSP4 | signal transduction | 11 | 5 |
| | NR4A3 | transcriptional activation | 3 | 12 |
| | SOD2 | oxidative stress | 3 | 11 |
| | STAT6 | signal transduction | 0 | 10 |
| | TBC1D4 | glucose homeostasis | 3 | 9 |
| | PPRC1 | mitochondrial biogenesis | | 9 |

### Supplementary Table 4. Network regulators involved in Rett Syndrome

| Regulator | Functions & Pathways | References |
|---|---|---|
| BDNF | • Involved in neuronal development, synaptic transmission, and plasticity<br>• Declines with the onset of Rett-like neuropathological and behavioral phenotypes | Li et al. 2014 |
| FCRL2 | • Involved in neuronal development, synaptic transmission, and plasticity<br>• Regulated by MeCP2 & implicated in Fragile X | |
| FMR1 | • Regulated by MeCP2 & implicated in Fragile X syndrome | Coffee et al. 1999; Liu et al. 2018 |
| MeCP2 | • Rett is caused by sporadic or germline mutations in MeCP2 on the X chromosome | |
| NTRK2 | • Involved in RNA granule formation in neurons<br>• Regulates MeCP2 | Abuhatzira et al. 2007 |
| PER1 | • Involved in the feedback loop of the core circadian clock<br>• MeCP2 binds and transcriptionally activates the circadian clock genes | Tsuchiya et al. 2015<br>Li et al. 2015 |
| PER2 | • Involved in the feedback loop of the core circadian clock<br>• MeCP2 binds and transcriptionally activates the circadian clock genes<br>• The oscillatory *Per2* gene expression is reduced in amplitude, which may contribute to sleep/wake cycle-related Rett symptoms | Tsuchiya et al. 2015<br>Li et al. 2015 |
| PUM1 | • Regulates MeCP2 by destabilizing MeCP2 transcripts | Rodrigues et al. 2016 |

**Supplementary Table 5: Top 30 compounds predicted by nemoCAD to reverse the Rett syndrome-like state of MeCP2 knockdown tadpoles to a healthy state.**
The corresponding prediction score, analogous to a distance from a healthy target state, is shown on the right. Lower score indicates better predicted efficacy. Bolded names indicate drugs that were tested.

| Drug | Score |
|---|---|
| **tyrphostin-AG-1478** | **20.5** |
| **sirolimus** | **28.875** |
| **vorinostat** | **33.625** |
| panobinostat | 37.875 |
| entinostat | 42.75 |
| trichostatin-a | 49 |
| palbociclib | 56.125 |
| nefazodone | 57.75 |
| pyrazolanthrone | 60.875 |
| tamoxifen | 65.75 |
| wortmannin | 73 |
| azacitidine | 76.625 |
| indole-3-carbinol | 77.375 |
| thapsigargin | 80 |
| diphenyleneiodonium | 83.5 |
| parthenolide | 89 |
| artesunate | 91.5 |
| chlorpromazine | 93.625 |
| pifithrin | 98 |
| **ivermectin** | **102.25** |
| cycloheximide | 102.375 |
| fluoxetine | 102.5 |
| **clozapine** | **103.375** |
| ethambutol | 104.75 |
| selamectin | 110.625 |
| triclosan | 115.375 |
| tunicamycin | 117 |
| mifepristone | 119.875 |
| genistein | 126.625 |
| cinobufagin | 126.875 |

**Supplementary Table 6: Protein targets shared between vorinostat and ivermectin identified within each of the three body regions using thermal proteome profiling.**

| Tail | Abdomen | Head |
|---|---|---|
| TRRAP | C9orf64 | si:ch211-222l21.1 |
| PLTP | slc5a6a | Prss2 |
| Hist1h3b | ATP5F1C | Acss3 |
| Rps8 | Acss3 | atp5fa1 |
| atp5fa1 | DIAPH1 | |
| noc4l | Ckap4 | |
| Rps6 | atp5fa1 | |
| | olfcg1 | |

**Supplementary Table 7. Pathway classes significantly enriched for Fragile X**

| Pathway Classes |
|---|
| Oxidative Stress |
| Phenylacetate degradation |
| Methylmalonyl pathway |
| 5-Hydroxytryptamine biosynthesis |
| Pantothenate biosynthesis |
| Leucine biosynthesis |
| Alanine biosynthesis |
| Nicotinic acetylcholine receptor signaling pathway |

**Supplementary Table 8. Pathways with the maximum number of gene matches**

| Pathways | Number of genes |
|---|---|
| Metabotropic glutamate receptor group III pathway | 5 |
| Alzheimer disease presenilin pathway | 4 |

| |
|---|
| CCKR signaling map |
| Dopamine receptor mediated signaling pathway |
| Endothelin signaling pathway |
| Gonadotropin-releasing hormone receptor pathway |
| Integrin signaling pathway |
| P53 pathway |
| Heterotrimeric G-protein signaling pathway- Gi alpha and Gs alpha mediated pathway |
| Inflammation mediated by chemokine and cytokine signaling pathway |

**Supplementary Table 9. sgRNA sequences used to generate tadpole models of Rett syndrome using CRISPR**

| Name | Spacer sequence | CHOPCHOP Rank | Location on MeCP2 |
|---|---|---|---|
| Xl-mecp2s-g01 | UCCAGAAAGCAGAAGCACAG | 1 | exon 3 |
| Xl-mecp2s-g02 | CGGUCUGUUAUUAGAGACAG | 2 | exon 2 |
| Xl-mecp2s-g03 | AUUUUGACACUUAAGUCCUG | 3 | exon 3 |
| Xl-mecp2s-g04 | CUUCUGCCUCUCCCAAACAG | 4 | exon 2 |
| Xl-mecp2l-g05 | AGGUCUGUUAUCAGAGACAG | 1 | exon 2 |
| Xl-mecp2l-g06 | UCGGCAUCUCCUAAAAAGAG | 2 | exon 3 |
| Xl-mecp2l-g07 | ACUUUAACACUUAAGUCCUG | 3 | exon 3 |
| Xl-mecp2l-g08 | CGGCAUCUCCUAAAAAGAGG | 4 | exon 3 |

### Supplementary Table 10. Primer sequences used for IDAA

| Name | Sequence |
|---|---|
| idaa-mecp2s.1-fw | AGCTGACCGGCAGCAAAATTGcttttacgaccgcctcttttaa |
| idaa-mecp2s.1-rv | cttctgtatcagggagaggaag |
| idaa-mecp2s.2-fw | AGCTGACCGGCAGCAAAATTGctaactctttctgtcaccgttg |
| idaa-mecp2s.2-rv | aaaaattaaaactggcttgccg |
| idaa-mecp2s.4.fw | AGCTGACCGGCAGCAAAATTGaacgaccagattttctttgctt |
| idaa-mecp2s.4.rv | atgatctgcttttctgttccac |
| idaa.mecp2L.1-fw | AGCTGACCGGCAGCAAAATTGattaaagctgtcttgccgaaaa |
| idaa.mecp2L.1-rv | ctaactctttctgtcaccgttg |
| idaa-mecp2L.2-fw | AGCTGACCGGCAGCAAAATTGctctttcatttccacagcccta |
| idaa-mecp2L.2-rv | aaagctttcctggactcttctc |
| idaa-mecp2L.3-fw | AGCTGACCGGCAGCAAAATTGtggataatgggttttcctggat |
| idaa-mecp2L.3-rv | gtctctgataacagacctcctc |
| idaa-FamFwd | /56-FAM/AGCTGACCGGCAGCAAAATTG |
